# Assessing phylogenetic information content and redundancy in hominin craniodental traits

**DOI:** 10.1101/2024.10.31.616875

**Authors:** Levi Yoder Raskin, Maja Šešelj, Bárbara Domingues Bitarello

## Abstract

Paleoanthropological phylogenetic inference is based on characters assumed to be phylogenetically informative and independent. Yet, our understanding of whether these criteria are met in published character supermatrices is limited. We assess the phylogenetic information content (PHIC) of 107 discrete craniodental traits from a widely-used hominin character matrix. We compare test topologies — inferred by permuting single traits or removing single traits, anatomical units (AUs), and operational taxonomic units — to the baseline topology, inferred from the unmodified matrix. In this dataset, only 31 traits have some degree of PHIC: 23 uniquely informative traits — sufficient, as a set, to closely approach the baseline topology — and eight redundant traits. No single AU, nor the combination of mandibular and dentition AUs, contains sufficient PHIC to approach the baseline topology, and only the maxilla contains more PHIC than expected. Therefore, phylogenetic placements of fossil hominins represented by isolated AUs should be regarded as putative until better-preserved specimens and more informative traits can be incorporated. Given the ubiquity of discrete morphological data in paleontology and that most of the history of life on Earth was only recorded through fossils, our methods should be broadly applicable to phylogenetic inference involving other paleontological clades.

## Background

Paleontological phylogenetics endeavors to reconstruct evolutionary relationships in fossil taxa primarily using morphological data, as ancient DNA is not available for most fossil taxa. The absence of genomic data imposes numerous challenges and introduces uncertainty in our ability to accurately reconstruct evolutionary relationships. Hominin phylogenetics is limited by the sparse nature of the fossil record, further complicated by differing rates of evolution across different anatomical structures [1], and homoplasy is potentially widespread [2–6]. However, because the hominin phylogenetic tree is inferred from said fossil record, understanding the relationships between those morphologies and the inferred tree will inherently be biased [5,7,8]. A way of circumventing this circularity is by adopting a tree-blind approach, i.e., relaxing the assumptions about the inferred tree being the correct one and instead understanding the impact that individual traits have on said inference.

Several hominin morphological traits demonstrate large amounts of conflicting phylogenetic signals, whereby different traits suggest different phylogenetic trees [6,9]. Some of this is likely due to homoplasy of masticatory characters [6]. For example, *Paranthropus robustus, P. boisei*, and *P. aethiopicus* appear to exhibit shared, derived craniodental features related to increasing bite force [3,10], but those characteristics may have arisen independently in two geographic regions in response to selection for increased bite force [2,3,5]. Our understanding of the relationship between the East and South African *Paranthropus (P. boisei* and *P. aethiopicus vs. P. robustus*, respectively) is further complicated by the morphology of *Australopithecus africanus*, also from South Africa. *P. robustus* may be more closely related to *Au. africanus* to the exclusion of the East African *Paranthropus* [3], but this interpretation contradicts phylogenetic analyses that favor *Paranthropus* as a monophyletic group [11–15]. Furthermore, the entirety of the hominin skull may exhibit high levels of homoplasy [2], as well as conflicting phylogenetic signals that may be partially due to homoplasy [6].

Another challenge regarding phylogenetic inference of fossil specimens is the mosaic nature of morphological evolution, observed in numerous hominins, including, for example, *H. sapiens*, and two South African fossil hominins with a seemingly confounding array of ancestral and derived characteristics relative to *Homo: Au. sediba* [16] and *H. naledi* [17]. Given that phylogenies built using parsimony rely on character similarities, placing such specimens in the hominin tree is extraordinarily challenging. Fundamentally, mosaic evolution implies a modular organization of the organism with varying rate and direction of change acting on those units [1]. Thus, understanding the influence of anatomical units and individual characters on the resulting phylogenetic hypotheses is critical to understanding mosaicism’s effect on parsimony-based phylogenetic inference.

Here, we use the maximum parsimony framework to we assess the relative contribution of each trait and anatomical unit in a well-established hominin character matrix of 107 craniodental traits [14] to the resulting tree topology and focus on how individual traits, units, and taxa affect phylogenetic placement. We use multiple approaches involving removal and permutation of individual traits and anatomical units from the phylogenetic inference to assess the uniqueness and phylogenetic information contained within these traits. We find that a small proportion of the traits effectively contribute phylogenetic information to the inferred topology and that isolated anatomical units lack enough information for robust phylogenetic inference.

## Results

### The baseline topology

The dataset compiled by Mongle et al. [14] consists of 20 operational taxonomic units (OTUs) (SOM Tables S1-S3), including 13 hominin fossil taxa, modern humans, and six non-human extant catarrhine primates, and limited by the amount of missing data in the fossil record. *K. platyops, Au. garhi* and *Au. anamensis* are particularly affected (79, 78, and 75 missing characters, respectively; Figure 1, SOM Table S2). Only three of the 107 traits have data from all extinct hominin OTUs: reduced canines (trait 51), molar crown area (trait 54), and relative enamel thickness (trait 59) (Figure 1, SOM Table S3). Because this dataset — and the hominin record in general — are sparse, understanding which characters are redundant or altogether devoid of information may help maximize the amount of relevant information gained from a scant fossil record.

**Figure 1.**
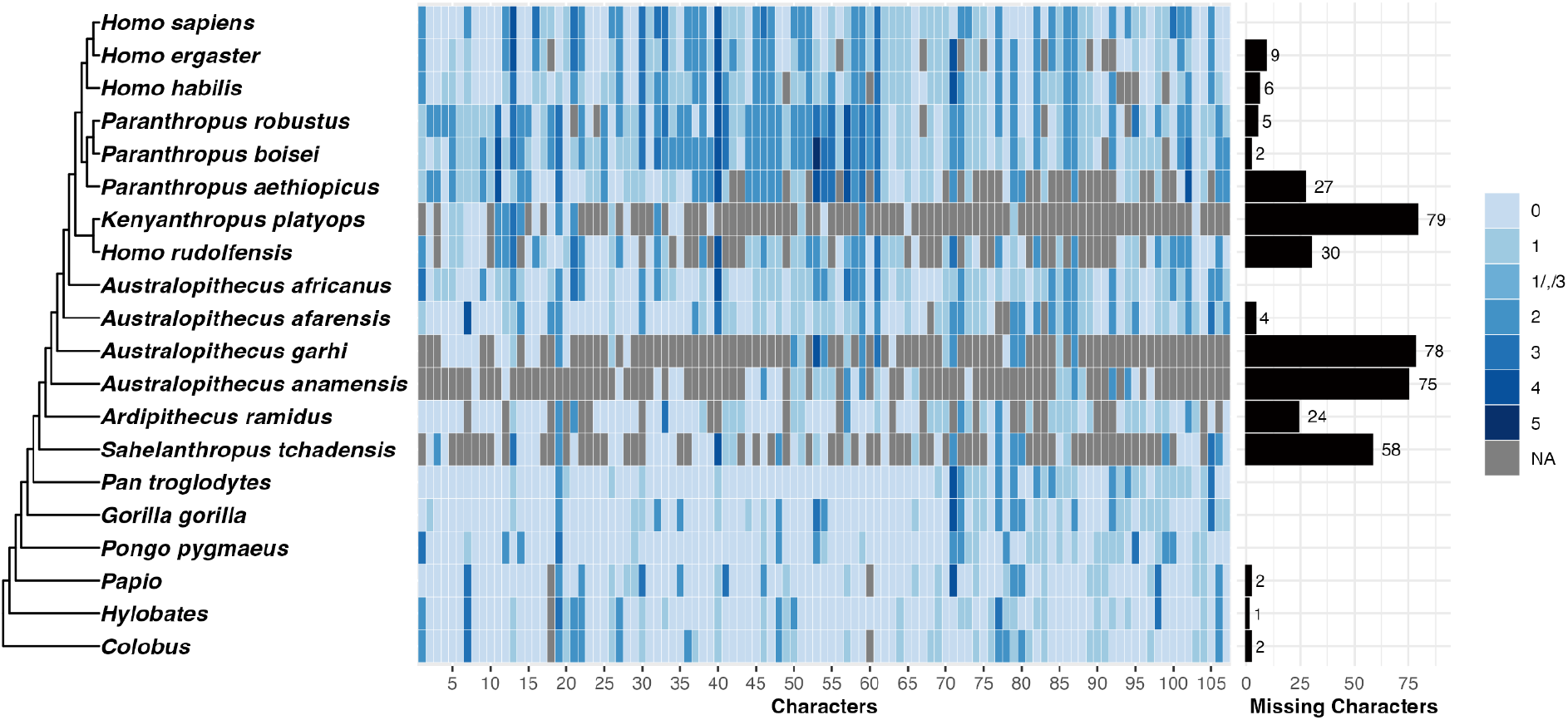
Baseline topology for 20 operational taxonomic units (OTUs). Left, the consensus tree topology (cladogram) inferred using branch and bound with maximum clade credibility. Center, coding for each taxon (rows) and character (columns). Character index numbers (1-107) and codings (blue tones; grey for missing data) correspond to those in SOM Table S1. Right, missing characters per OTU. The topology is rooted with *Colobus*.

Using all OTUs and traits, we inferred three maximum parsimony (MP) topologies (parsimony score 386) using the branch and bound algorithm (BAB, see Methods) and their consensus tree, hereafter referred to as the baseline topology (Figure 1). The baseline hominin + *Pan* subtree topology is broadly similar (Robinson Foulds distance *RF* = 6, see Methods) to Mongle et al.’s [14] parsimony-inferred topology (score 390), with two differences. First, our baseline topology grouped *H. rudolfensis* and *K. platyops* as sister taxa forming a sister clade to the *Homo*-*Paranthropus* clade (excluding *H. rudolfensis*), whereas Mongle et al.’s [14] places *K. platyops* as the outgroup to a monophyletic *Homo* with *Paranthropus* as a sister clade to *K. platyops + Homo*. Secondly, our baseline topology places *Au. garhi* as basal to *Au. afarensis*, whereas in Mongle et al.’s [14] *Au. afarensis* is basal to *Au. garhi*.

Our goal here was not to infer the putatively “most correct” hominin phylogeny. Rather, we explored how different hominin craniodental characters affect the inferred topology. Our baseline topology represents a reasonable hypothesis against which all other test topologies are compared and thus affects the magnitude of *RRF* and 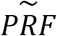. Using Mongle et al., [14]‘s parsimony-inferred topology as a baseline, values were generally higher, as expected, but highly correlated with those from our baseline topology (*p* < 0. 001 in both cases, Kendall’s τ correlation, SOM Fig. S1).

### Cladistic information content (CIC)

One of our key goals was to characterize the degree of phylogenetic information content (PHIC) of individual traits. Here, we make an explicit distinction between CIC and PHIC. CIC is a function of the number of states and how many OTUs carry each state, but does not consider topology [18]. In contrast, our approaches (Table 1 and Methods) attempt to capture the PHIC relative to the proposed phylogeny, and as such are conditioned on the tree topology. Throughout, we use CIC as a measure of the “potential” of a trait to harbor phylogenetic information; thus, a trait with *CIC* = 0 should not be expected to have any PHIC (as measured by our approaches), whereas a trait with relatively high *CIC* has the potential to contain some degree of PHIC.

**Table 1.** Summary of analyses to quantify the phylogenetic information content (PHIC) and uniqueness of each character or anatomical unit. RF: Robinson-Foulds distance; AU: anatomical unit; ^a^, following (Cotton & Wilkinson, 2008); ^b^, higher and lower here are relative to other traits (CIC, PRF, RRF, TRRF), units (URRF, SURF), or to a null distribution (URRF, SURF); ^d^, the baseline topology (Figure 1).

| Name | Metric | Interpretation <sup>b</sup> |
| --- | --- | --- |
| Cladistic Information Content (CIC) | $CIC = -\ln(\hat{p})$, where $\hat{p}$ is the proportion of binary trees for which the trait is homoplasy free.<sup>a</sup> | The *a priori* grouping potential of a trait based on the number of and distribution of states across taxa, regardless of a trait's fit to a tree. Higher values indicate higher potential. |
| Permutation RF distribution (PRF) | $\tilde{PRF}$: the median of the PRF distribution of $RF$ values between the baseline and each permutation test topology. | The degree of PHIC contributed by a trait to the baseline<sup>d</sup>.<br>Lower $\tilde{PRF}$: lower PHIC.<br>Lower $\tilde{PRF}_{IQR}$: lower instability (variance) |
| Trait removal RF (RRF) | $RRF$: The RF between the baseline and a test topology inferred from a character matrix with one trait removed. | The degree of unique PHIC contributed by a trait to the baseline.<br>Lower $RRF$: less PHIC overall or redundant PHIC. |
| Unit removal RF (URRF) | $URRF$: The RF between the baseline and a test topology inferred from a character matrix with all traits in a given AU removed. | The degree of unique PHIC contributed by an AU to the baseline.<br>Lower $URRF$: lower PHIC overall or redundant PHIC. |
| Single unit RF (SURF) | $SURF$: RF between the baseline and a test topology inferred solely from characters in a given AU. | The PHIC in an anatomical unit in relation to size-matched random set of traits.<br>Lower $SURF$: higher PHIC than expected. |
| Taxon removal RF (TRRF) | $TRRF$: RF between the baseline and a test topology inferred missing each taxon one at a time. | The PHIC contributed by each OTU to the baseline.<br>Lower $TRRF$: lower PHIC. |

The *CIC* values across the 107 traits have a mean of 12.673, standard deviation of 5.297 (Figure 2A), and ranging from 0 (character 83, “ossification of the petrous aspect”, coded as 0 in *H. sapiens* and 1 or “?” [missing] in all other OTUs) to 25.841 (character 54, “molar crown area”) (SOM Table S5). Because we are using *CIC* as a relative measure, hereafter we express *CIC* values in terms of their rank (*CIC*_*rank*_) based on their CIC value divided by the number of traits. The resulting quartiles can be directly interpreted: “low CIC” (*CIC*_*rank*_ < 0. 25), “mid CIC” (0. 25 ≤ *CIC*_*rank*_ < 0. 75), and “high CIC” (0. 75 ≤ *CIC*_*rank*_).

**Figure 2.**
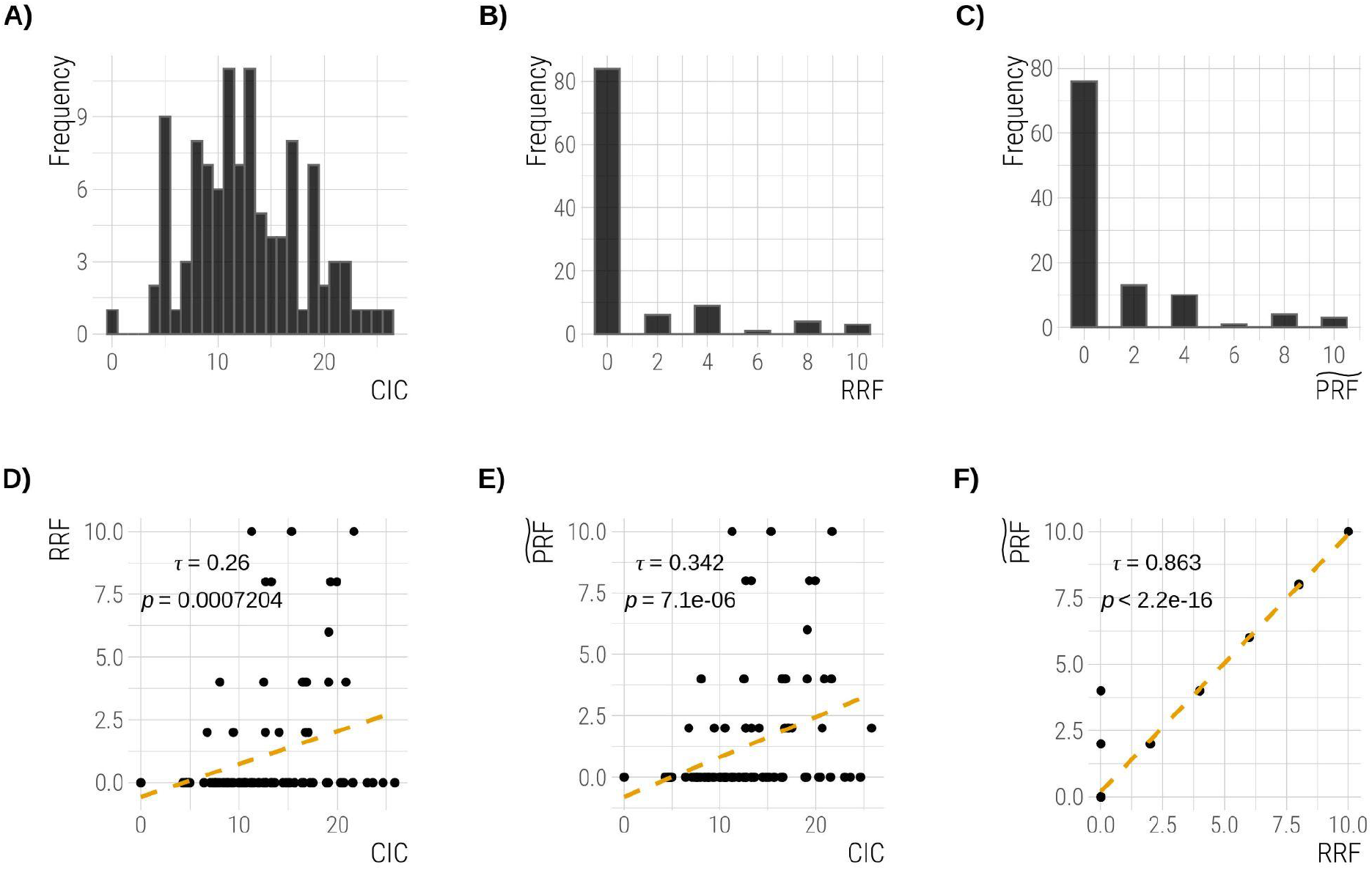
Distributions of different metrics and their correlations across 107 characters. A) cladistic information content, B) removal Robinson-Foulds distance, C) median permutation Robinson-Foulds distance (250 permutations for each trait), D) *RRF vs. CIC*, E) 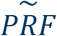 *vs. CIC*, and F) 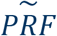 *vs. RRF*. D)-F) correlation coefficient (τ, Kendall’s rank correlation, regression line in orange) and *p*-values.

### Phylogenetic information content (PHIC)

We proposed two main approaches to quantify the degree of PHIC: RRF and PRF (Table 1). A common method to assess the relationship between traits and trees consists of permuting the tip values across OTUs [19,20]. This allows quantifying whether the observed distribution of traits along a tree is likely to have occurred randomly. We elaborate on this idea by testing how arrangements that contain no phylogenetic signal change the inferred topology: assuming that each trait in the dataset exhibits some amount of phylogenetic signal, does the disruption of that signal by removal or permutation result in changes to the inferred topology? We hypothesized that if a trait contributes PHIC to the inferred topology, the permutation of its values would result in a topological distance to the baseline topology, and that its removal would either result in an increased distance (if the information carried by the trait is unique) or not (if the information is redundant with information carried by a different trait that was not removed).

The distributions of both 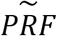 (median *PRF*) and *RRF* (Table 1) are right-skewed (Figure 2B-C), with same range (between 0 and 10), same medians (0 for both), and similar means (1.08 and 1.25 for *RRF* and 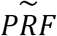, respectively). Across all traits, *RRF* and 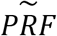 are highly correlated (τ = 0. 863, *p* < 2. 2 × 10^−16^, Figure 2F). This is sensible given that in one case (RRF) we remove a trait entirely, and in the other (PRF) we destroy the information it carries. We only inferred even-numbered *RRF* and 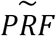 values (Figure 2B-C) — we never inferred a polytomy in the hominin ingroup. Both metrics are also modestly but significantly positively correlated with *CIC* (*CIC vs. RRF*: τ = 0. 26, *p* = 0. 00072; *CIC vs*. 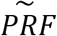, τ = 0. 342, *p* = 7. 1 × 10^−6^, Figure 2D-E). Based on our criteria for classifying traits as informative/uninformative and unique/redundant (Figure 3A), 76 traits (71%) were deemed uninformative because permutation across OTUs has no effect on the baseline topology. The remaining 31 traits (29%) contain variable degrees of redundant or unique PHIC: 23 (21.5 % of all traits) are non-redundant (“unique”) in the PHIC they carry (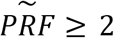 and *RRF* ≥ 2), and the remaining eight traits, while potentially informative 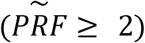 carry variation that is captured by other traits — i.e, they are redundant (*RRF* = 0). Throughout the results and discussion, we often highlight traits that are commonly studied in paleoanthropology.

**Figure 3.**
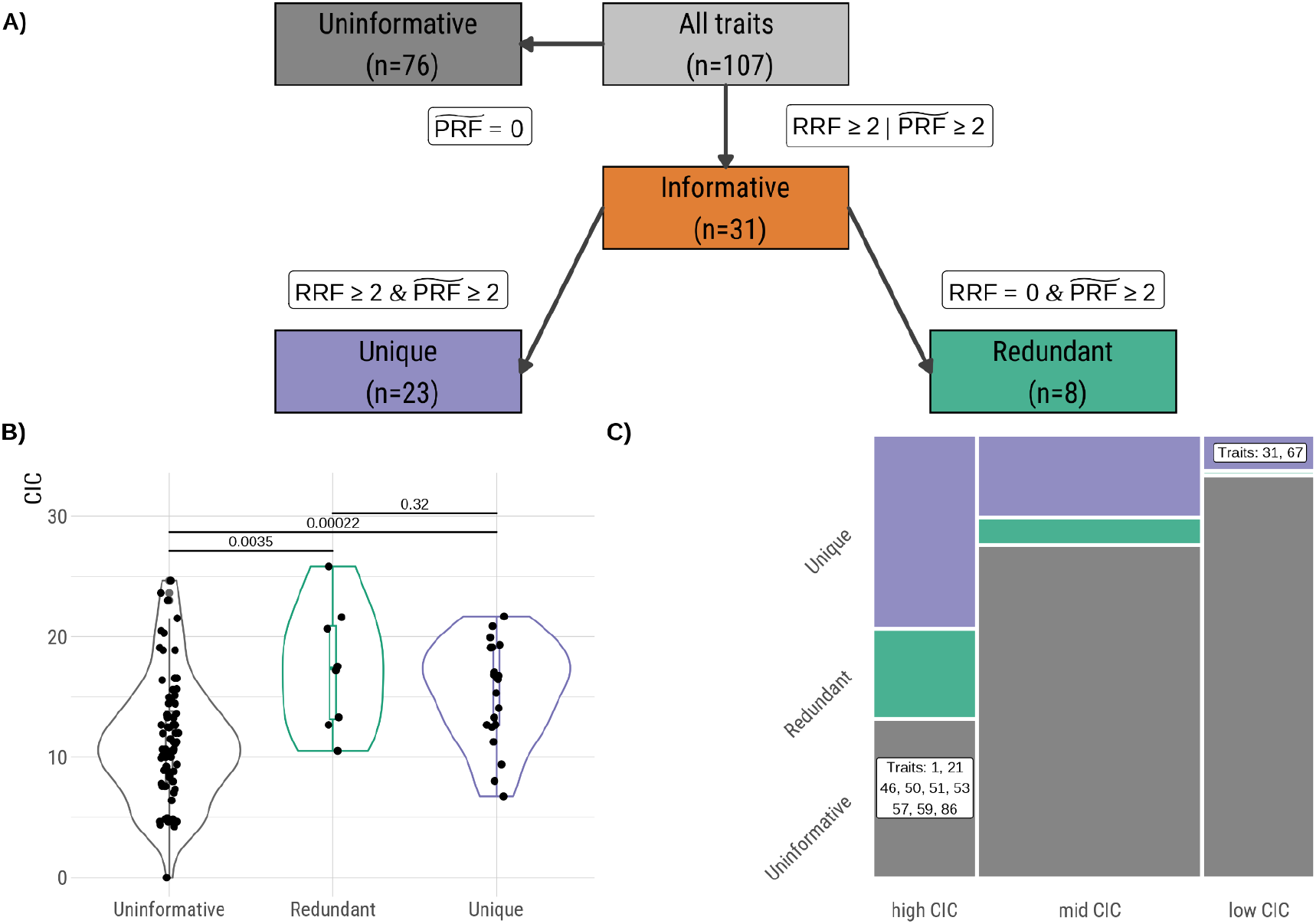
Classification of traits in terms of phylogenetic information content. A) A schematic explaining the categorization of the 107 traits based on *RRF* and 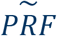. B) *CIC* distributions across uninformative and informative (redundant or unique) traits. Boxplots are defined by the 25% and 75% quantiles, and whiskers reach at most ± 1. 5 × *IQR. IQR*, the interquartile range. Horizontal bars show comparisons and *p*-values (MWU). C) Proportions of unique, redundant, and uninformative traits within bins of *CIC* (low, mid, high, see Methods). Traits annotated in the figure are those that fall within the “uninformative” and “high CIC”(bottom right) and “unique” and “low CIC” (top right) intersections. Trait names and definitions are listed in SOM Table S1.

Confirming our intuition that *CIC* values are a measure of the “potential” of a trait to harbor phylogenetic information, we find that the only trait with *CIC* = 0 (character 83) also has *RRF* = 0 and 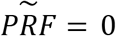 and can be safely assumed to be devoid of PHIC (SOM Table S6). As expected, *CIC* is strongly negatively correlated with the number of OTUs with missing data (τ =− 0. 412, *p* = 3. 494 × 10^−9^), whereas *RRF* and 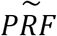 are similarly but less intensely affected (*RRF*: τ =− 0. 174, 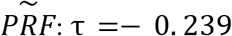, *p* = 0. 0029). The negative correlation persists for *CIC* when looking only at the 31 informative traits (τ =− 0. 334, *p* = 0. 014), whereas for *RRF* and 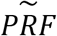 it disappears (*RRF*: τ = 0. 122, 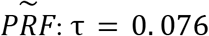, *p* = 0. 614).

Lastly, *CIC* is generally higher for informative traits than for uninformative ones (MWU, uninformative *vs*. redundant, *p* = 0. 0035; uninformative *vs*. redundant: *p* = 0. 00022, Figure 3B). Nevertheless, no significant difference was detected between *CIC* values for unique *vs*. redundant traits (MWU, *p* = 0. 32, Figure 3B), indicating that *CIC* alone would not have allowed this level of insight — uniquely *vs*. redundantly informative — about the traits.

#### Uniquely informative traits

The 23 uniquely informative traits (SOM Table S7) are those whose removal and permutation both affect the baseline topology — which does not entail they are intrinsically “better” or informing the true topology (see Discussion). For this set of traits, *RRF* and 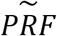 are perfectly correlated, so we mention only their 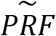 values. Their *CIC*_*rank*_ values span from 0.131 (character 67) to 0.963 (character 33, Figures 2A, 3B, SOM Table S5), once again indicating that *CIC* alone would not have allowed the detection of these traits as a set.

The three most informative traits (SOM Table S7) are the size of the postglenoid process (character 33, *CIC*_*rank*_ = 0. 963, 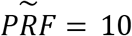, *PRF*_*IQR*_ = 6, see Table 1 and Methods), the orientation of the mandibular premolar row arcade shape (character 62, *CIC*_*rank*_ = 0. 701, 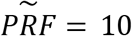, 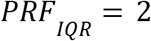, *PRF*_*IQR*_ = 2), and the prominence of the lingual ridge of the mandibular canine (character 52, *CIC*_*rank*_ = 0. 430, 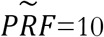, *PRF*_*IQR*_ = 0). Their individual removal or permutation results in major changes to the baseline topology, in particular affecting the position of the *Paranthropus* clade (SOM Fig. S3-S5). For example, in both the removal and the most commonly inferred permutation topology for character 33 (SOM Fig. S3), in contrast to the baseline, *Homo* is monophyletic and the *Paranthropus* clade is inferred as differentiating much earlier, with *Paranthropus* as an outgroup to the clade containing *Au. garhi, Au. afarensis, K. platyops, Au. africanus*, and *Homo*.

To ensure these 23 uniquely informative traits were not redundant to one another and not all informative to a few, tenuously supported relationships in the baseline topology (as traits 33, 52, and 62 and their effect on the *Paranthropus* clade), we compared RRF test topologies from the 23 uniquely informative traits to each other: these ranged from *RF* = 0 to *RF* = 12, with most topologies being quite distant from one another with 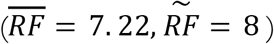 (SOM Fig. S6).

To gain further insight into our *RRF* and 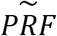, we argue that if a trait is uniquely informative but falls within the low *CIC* bin, it provides more phylogenetic information than its potential based on *CIC* alone. We find two traits in this category (Figure 3C, top right): the horizontal distance between the temporomandibular joint and the M^2^/M^3^ (character 31, *CIC*_*rank*_ = 0. 206, 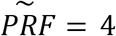) and the configuration of the superior orbital fissure (character 67, *CIC*_*rank*_ 0. 131, 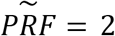). Character 67’s low *CIC* can be explained by its 8 missing OTUs and only 2 coded states (Figure 1, SOM Table S3). Nevertheless, it is highly informative to the baseline topology: removal of this trait places *K. platyops* as a sister group to *Homo + Paranthropus*, in contrast to the baseline topology, where *K. platyops* is grouped with *Homo + Paranthropus*. Conversely, if a trait is uninformative and falls within the high CIC bin we argue that the trait provides less information than expected based on *CIC*.

#### Redundant traits

Among the 31 informative traits, we found eight traits which, though informative 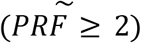, contributed redundant information (*RRF* = 0, Figure 3A). Among these, five have high *CIC*, including facial prognathism (character 13, *CIC*_*rank*_ = 0. 925) and molar crown area (character 54, *CIC*_*rank*_ = 1). Despite having the highest *CIC* across all traits, the latter contributes no unique PHIC (*RRF* = 0, 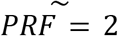, *PRF*_*IQR*_ = 4), i.e, it is redundant with at least one other character in the matrix.

To test the ability of these uniquely informative and informative traits to resolve the baseline hominin topology (Figure 1) as a set, we inferred two test topologies: one from the 31 informative traits and one from the 23 uniquely informative traits. The one inferred from all 31 informative traits differed from the baseline topology by *RF* = 12 and is not significantly different from the null expectation for this number of traits (*p* = 0. 58), but the test topology inferred from just the 23 uniquely informative traits differed from the baseline topology by only *RF* = 4 and is significantly less distant than expected (*p* = 0. 02; SOM Fig. S7).

#### Uninformative traits

More than 70% of the traits are, individually, uninformative (Figure 3A). Though the criterion to group them as such was that 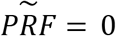, we note that all of them also have *RRF* = 0 (SOM Table S5). Among these 76 uninformative traits, nine are also high CIC traits (Figure 3C, bottom left), once again showing that CIC alone does not predict whether a trait will actually inform an inferred tree topology, but rather of a trait’s potential to have such an effect. These include widely studied traits in paleoanthropology: the reduction of incisors (character 50, *CIC*_*rank*_ = 0. 972, *PRF*_*IQR*_ = 4) and the reduction of canines (character 51, *CIC*_*rank*_ = 0. 981, *PRF*_*IQR*_ = 2).

Furthermore, 57 uninformative traits also had *PRF*_*IQR*_ = 0 — a majority of topologies built from random permutations of their values did not differ from the baseline. Among these 57 are traits associated with *Paranthropus*: sagittal cresting (character 20, *CIC*_*rank*_ = 0. 589, *PRF*_*IQR*_ = 0), anterior pillars (character 4, *CIC*_*rank*_ = 0. 112, *PRF*_*IQR*_ = 0), reduction of incisors (character 50, *CIC*_*rank*_ = 0. 972, *PRF*_*IQR*_ = 4), and relative enamel thickness (character 59, *CIC*_*rank*_ = 0. 944, *PRF*_*IQR*_ = 0). Strikingly, for 22 of these 57 (SOM Table S6), *RF* = 0 for all permutation test trees — none of the 250 permutations resulted in a topology different from the baseline. Included among them are the inclination of the nuchal plane, distinctiveness of the angular process of the mandible, and robusticity of canines (SOM Tables S5-S6). Unexpectedly, *K. platyops* is not coded for any of these 22 traits (*p* = 0. 00045, from 100,000 resampled sets of 22 traits).

To test that these 22 “highly uninformative” traits had no impact on the baseline topology, we inferred a test topology excluding these 22 traits from the dataset. Interestingly, we found that it was *RF* = 4 from the baseline, showcasing that while our approach identifies uninformative traits on their own, these categorizations may not smoothly transition to identifying blocks of traits.

### Effects of anatomical units on the inferred topology

We assigned each of the 107 craniodental traits to one of 10 anatomical units (AUs), where nine represent individual bones and the tenth (the “complex” AU) encompasses traits that include more than one bone (Table 2). The number of traits per unit ranges from *n* = 1 (nasal) to *n* = 25 (dentition). To gain further insight on the amount of information contained within individual AUs relative to the baseline topology, we implemented two approaches. In one approach (URRF) we looked at distances of test topologies obtained by removing an entire AU to the baseline, and in the other (SURF) by only including traits in those AUs in the test topology inference (Table 1). Because the AUs contain different numbers of traits, we contextualize the SURF and URRF values to their respective null distributions, obtained by sampling “pseudo AUs” of the same size as actual AUs (see Methods).

**Table 2.** Robinson-Foulds distances (RF) of each region of the skull. AU, anatomical unit. URRF, unit removal RF; SURF, single unit RF. *p*-values based on sampling 100 pseudo-AUs of same length as the AUs (SOM Table S1 and Methods). Nasal (*n* = 1) is omitted (see Methods). ^a^, the number of MP trees (BAB algorithm). ^b^ prohibitive runtime (see Methods). Trait names and definitions are listed in SOM Table S1.

| AU<br>(number of traits) | Traits | URRF | SURF | MP<br>Trees<sup>a</sup> |
| --- | --- | --- | --- | --- |
| Zygomatic<br>(n=7) | 9, 10, 11, 77, 81, 102, 107 | 10 (*p* = 0.26) | 22 (*p* = 0.19) | 2,350,575 |
| Dentition<br>(n=25) | 49, 50, 51, 52, 53, 54, 55, 56, 57,<br>58, 59, 60, 61, 70, 84, 85, 86, 87,<br>88, 89, 90, 91, 92, 97, 99 | 14 (*p* = 0.23) | 20 (*p* = 0.36) | 1,278 |
| Frontal<br>(n=9) | 19, 26, 67, 71, 75, 78, 79, 80, 105 | 4 (*p* = 1.00) | 20 (*p* = 0.74) | 356,440 |
| Occipital<br>(n=9) | 18, 21, 30, 39, 40, 42, 68, 96, 106 | 8 (*p* = 0.60) | 20 (*p* = 0.74) | 46,167 |
| Maxilla<br>(n=13) | 2, 3, 4, 5, 6, 7, 8, 12, 13, 15, 76, 82,<br>101 | 12 (*p* = 0.03) | 18 (*p* = 1.00) | 37,014 |
| Temporal<br>(n=14) | 24, 25, 27, 29, 33, 34, 35, 36, 37,<br>38, 65, 69, 72, 83 | 8 (*p* = 0.93) | 14 (*p* = 0.31) | 9,278,325 |
| Mandible<br>(n=15) | 32, 43, 44, 45, 46, 47, 48, 62, 66,<br>73, 74, 93, 94, 95, 98 | 10 (*p* = 0.45) | 12 (*p* = 0.15) | 72,520 |
| Complex<br>(n=9) | 14, 16, 17, 22, 28, 31, 41, 103, 104 | 8 (*p* = 0.60) | —<sup>b</sup> | —<sup>b</sup> |
| Parietal<br>(n=5) | 20, 23, 63, 64, 100 | 0 (*p* = 0.60) | —<sup>b</sup> | —<sup>b</sup> |

URRF is significantly negatively correlated with the number of traits on each unit (τ =− 0. 554, *p* = 0. 0496, Kendall). Additionally, while in an absolute sense we find different URRF values associated with the different AUs, with dentition containing the most phylogenetic information (*URRF* = 14) followed by the maxilla (*URRF* = 12), mandible (*URRF* = 10), and zygomatic (*URRF* = 10, Table 2), only the maxilla has a significantly higher URRF than expected given the number of traits in the AU (*p* = 0. 03, *p* − *adj* = 0. 27, Table 2, SOM Fig. S8). However, the topology inferred solely from maxillary traits is very distant from the baseline one and not unexpected (*SURF* = 18, *p* = 1. 00). All SURF values fall well within the expected range given their respective null distributions. However, SURF values are marginally correlated with the number of traits in each AU (τ =− 0. 632, *p* = 0. 0572, Kendall’).

Though the phylogenetic topologies inferred solely from the mandibular and dentition AUs are well within expectation given the number of traits they contain (Table 2, SOM Fig. S8), we nevertheless consider them in greater detail given the prevalence of isolated mandibles and teeth in the hominin fossil record. Both differ substantially from the baseline topology (Figure 4A-B). Among other differences, in the mandibular-traits-only topology (*SURF* = 12, Figure 4A), *Ar. ramidus, Au. anamensis*, and *S. tchadensis* are not inferred in the hominin clade: *Ar. ramidus* is inferred as a sister taxon to *Pan troglodytes*, and *Au. anamensis* and *S. tchadensis* are inferred in a clade with *G. gorilla* and *Papio*. Similarly, in the dentition-only topology (*SURF* = 20, Figure 4B), all hominins are inferred in a monophyletic hominin clade, but *S. tchadensis* is inferred as more derived than *Ar. ramidus* and *Au. anamensis*, while *H. ergaster* is inferred as a sister group to a clade containing *H. sapiens, Au. garhi*, and *Paranthropus*.

**Figure 4.**
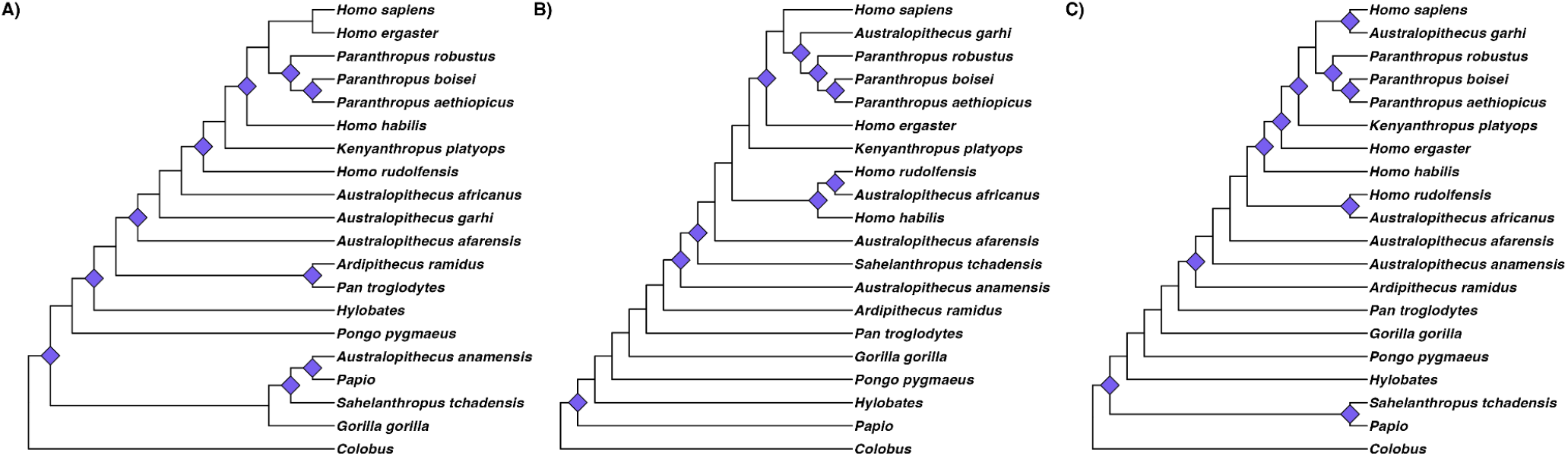
Hominin phylogenetic tree topologies inferred from sets of characters from different anatomical units. Consensus tree topologies inferred using the branch and bound algorithm with maximum clade credibility using A) solely mandibular characters (*n* = 15, *SURF* = 12), B), solely dental characters (*n* = 25, *SURF* = 20), or C) both dental and mandibular characters (*n* = 40, *RF* = 16). All topologies are rooted with *Colobus*. Purple diamonds indicate all changes in relation to the baseline topology, but only the differences in the hominin + *Pan* subtree topology were considered in the RF calculation.

While all reported topologies here are the consensus most parsimonious trees with the highest clade credibility (see Methods), the BAB algorithm returned an astounding number of MP trees for all SURF analyses — 72,520 and 1,278 equally parsimonious trees for the mandibular and dentition-only trees, respectively (Table 2). The number of MP tree topologies is not significantly correlated with the number of traits contained in the unit (τ =− 0. 390, *p* = 0. 224, Kendall).

Because mandibular and dental traits are so often used to place fossil hominin specimens, we also assessed how far from the baseline tree is a test tree inferred jointly from mandibular and dental traits (*n* = 40, Figure 4C). While using mandibular and dental traits jointly more effectively specifies tree space (only one MP tree is returned by BAB), the MP topology is substantially distant from the baseline (*RF* = 16) and not any better at resolving the baseline topology then random sets of 40 traits though this is not unexpected (*p* = 0. 53). Those units jointly lead to a topology that is more distant from the baseline than the mandible-only topology (Figure 4).

### Effects of each OTU on the inferred phylogenetic tree

Removing each OTU individually from the dataset (TRRF, Table 1) allowed an assessment of the relative contribution of each OTU to the inferred baseline topology (SOM Table S8). These varied from OTUs that do not affect the inferred hominin + *Pan* topology (*TRRF* = 0, *P. aethiopicus, Au. garhi, Ar. ramidus*, and *Po. pygmaeus*; SOM Table S8) to various degrees of effect, ranging from moderate (*TRRF* = 4: *Colobus, G*.*gorilla, Hylobates, H. ergaster, Papio, P. boisei, S. tchadensis*, SOM Figs. S18-S24; *TRRF* = 6: *Au. afarensis, Au. africanus, H. habilis*, SOM Figs. S15-S17) to high (*TRRF* = 8: *Au. anamensis, H. sapiens, H. rudolfensis, K. platyops, P. robustus*, SOM Figs. S10-S14; *TRRF* = 10: *Pan troglodyte*s, SOM Fig. S9). We found no correlation between *TRRF* and the number of missing traits in a given OTU (τ =− 0. 037, *p* = 0. 838). The consensus MP tree inferred when *K. platyops* is removed due to the long-standing questions surrounding the validity of this taxon. In contrast to the baseline, this topology (SOM Figure 12) positions *Homo* as monophyletic and *Au. garhi* and *Au. africanus* as sister taxa to the *Paranthropus* clade.

## Discussion

We applied several analytical approaches to an existing dataset of 107 hominin craniodental characters and 20 OTUs [14] to evaluate to what degree those characters contribute useful information for the phylogenetic placement of fossil hominins. We compared all test topologies against a baseline —the best-supported MP topology inferred from the full character matrix. Our approaches do not require knowing the most accurate (genomic) phylogeny and, accordingly, we make no claims about the baseline topology being closest to the truth. In fact, given our criteria, it is entirely possible for a trait to be uniquely informative while informing an incorrect phylogeny, or that it informs particular placements that are otherwise (in its absence) not robust. Rather, we report that the character matrix used here — and widely across hominin phylogenetic studies [6,11–15,21] — includes traits and taxa that, at the present time, may not be meaningful and perhaps confounding, to the understanding of hominin evolution. Throughout this discussion, we contextualize the relevance of our findings for paleoanthropology.

### Uniquely informative traits

The baseline topology is dependent on a small proportion of the traits, with 31 informative characters (∼29%), 23 of which are non-redundant. Our test topology with the 23 uniquely informative traits was very similar to the baseline (*RF* = 4), indicative that our approach is well-suited to identifying the majority of uniquely informative traits in a dataset though also conservative in that these two trees are not identical. However, the one inferred from the 31 informative traits was not. This is somewhat expected: the uniquely informative traits are, by definition, not redundant, so the addition of redundant traits adds new information.

The most phylogenetically informative traits were the size of the postglenoid process, the prominence of the lingual ridge of the mandibular canine, and the orientation of the mandibular premolar row dental arcade. Their observed information, however, when removed or permuted, comes from an inferred basal split of *Paranthropus*, a position that is inconsistent with our baseline topology and most of the literature [10,12–15,21–25]. If changes to one of these traits is enough to infer *Paranthropus* as a more basal hominin — a placement also found with a combination of temporal and morphological data [26] — then the usage of *Paranthropus* to contextualize the origins of *Homo* may need to be reevaluated. Our findings thus suggest that the derived position of *Paranthropus* often seen in the literature and our baseline topology is tenuously supported. However, two key traits in discussions regarding the monophyly of *Paranthropus* — anterior pillars and sagittal cresting in males [10,23–25] — are uninformative. Their lack of information further supports *Paranthropus* monophyly, as *Paranthropus* is still inferred as monophyletic (despite discussions otherwise, [2,3]) even without these supposedly critical characters.

### Uninformative and redundantly informative traits

We distinguish between a trait that carries no phylogenetic information towards the baseline topology and a trait that would otherwise be informative but is redundant. We found that 76 (∼71%) of the characters in the matrix are uninformative to the baseline topology, and eight (26% of the 31 phylogenetically informative traits) are redundantly informative. While one may be tempted to include as many traits as possible in fossil morphological phylogenetic inference, traits that are either uninformative or redundant should be treated with caution in systematic arguments. Redundancy in variation captured by multiple characters artificially inflates the weighting of those characters in a parsimony analysis [14,27], which our combination of RRF and PRF metrics can detect.

We detected informative and uninformative traits where a cladistic information content metric alone would have failed. Among the 14 high *CIC* traits that are either redundantly informative or uninformative, 50% are dental, including relative enamel thickness (uninformative) and molar crown area (redundant), suggesting caution in using dental traits in hominin phylogenetics. At the same time, a trait’s lack of information to the baseline topology does not necessarily mean they lack utility in all contexts. Rather, our findings suggest that specific tests are required to conclude in which taxonomic level or temporal context these traits may be informative.

The uninformative nature of the foramen magnum position is somewhat surprising given its intermediate *CIC* and that it is continually implicated in bipedal transitions [28–30]. The foramen magnum position, as operationalized in this dataset, may be informative across catarrhines but is not within hominins, reinforcing the importance of assessing which taxonomic levels are used to estimate phylogenetic signal and at which taxonomic levels are said estimates applied. The foramen magnum exemplifies issues with how continuous traits are discretized. While the hominin clade is defined by a progressively increasing capacity for bipedalism [31], the inevitable information loss from reducing continuous variation into discrete character states may result in the discretized trait no longer carrying enough information to impact inferred relationships. If this is true, then our finding may not indicate irrelevance of the foramen magnum, but rather insufficient information resulting from discretization of a continuous trait. Traits associated with bipedalism, like the foramen magnum position, may thus represent a strong argument for the incorporation of continuous traits into phylogenetics, as others have suggested [32,33].

Interestingly, our test topology without the 22 most uninformative traits was very similar but not identical to the baseline (*RF* = 4), indicating that even traits that singularly have no effect on the topology may carry information as part of a set, presumably due to the weight of characters in the parsimony analysis.

### Effects of anatomical units

Using the methods, dataset, and AU definitions used here, we find that isolated AUs carry low PHIC. Only the removal of the maxilla AU affected the tree more than expected for random sets of traits. The frontal bone’s lack of information is not surprising given that much of the potentially relevant variation occurs in Middle Pleistocene hominins not included here (and even in those OTUs it does not appear to be phylogenetically informative: [34,35]). In contrast, the lack of information in the parietal is surprising given the consistent increase in encephalization over hominin evolution and the correlation between parietal morphology and encephalization [36]. Relevant parietal information was possibly captured by non-parietal traits (e.g., cerebellar morphology). Additionally, all five traits on the parietal were uninformative. Thus, we cannot rule out that our findings simply reflect how we assigned traits to AUs and the specific composition of the dataset.

Individual AUs cannot discriminate across much of the tree space in isolation. First, we find a high number of resolved MP trees for all AUs. None of the AUs alone provides enough information to retrieve the baseline topology tree nor more information than expected for size-matched sets of random traits. These findings hold special relevance for all clades with fragmentary fossil records. For example, the tree topology from mandibular characters only (excluding mandibular dentition traits, grouped with “dentition”) was remarkably distant from the baseline, mostly affecting *Au. anamensis, S. tchadensis, Ar. ramidus, H. habilis*, and *H. rudolfensis*. Similarly, dental characters alone are insufficient to differentiate early *Homo* from *Australopithecus*, evidenced by *H. rudolfensis* and *H. habilis* being inferred as closer to *Au. africanus* than to any other *Homo*. Combining dental and mandibular characters does not lead to a greater than expected PHIC than expected for a random set of traits and the resulting MP topology places *H. rudolfensis* as a sister taxon to *Au. africanus* and *H. sapiens* as a sister taxon to *Au. garhi*. Isolated mandibular remains are relatively common in the hominin fossil record and much research has relied on mandibular morphology and characters to make systematic arguments [37–42]. However, our findings are consistent with past research warning against using solely mandibular traits [43,44] when inferring hominin phylogenetic relationships, and we extend that warning to mandibular fossils that preserve some dental traits.

We urge caution and skepticism when attempting to phylogenetically place fossil remains based on isolated AUs alone, at least on the basis of the characters included in the Mongle et al. [14] dataset, AUs as defined here, and the parsimony framework. Lastly, we defined AUs solely based on the bones of the skull. Future analyses may include a prior grouping based on known functional correlation across those structures [7].

### Effects of individual OTUs

We investigated the effect of each OTU on the inferred topology because the dataset we analyzed [14] includes taxa whose validity has been questioned [45,46]. Moreover, outgroup choices can affect ingroup topology. Lastly, the process of assigning fossils to OTUs cannot be independently confirmed with genomic methods, and thus may bias the inferred topology in poorly understood ways.

The removal of all outgroups except *Po. pygmaeus* results in different resolved hominin topologies, supporting the utility of including more outgroups in hominin phylogenetics [47]. *Pan* has the largest effect, supporting concerns that using only *Pan* as outgroup for hominins may increase bias from *Pan*-specific characters [48,49]. Among hominins, *K. platyops* has the highest TRRF (shared with *Au. anamensis, H. sapiens, H. rudolfensis* and *P. robustus*) despite having the largest number of missing traits. Its impact on the inferred topology is critical to understanding the origins and phylogenetic validity of our genus. When *K. platyops* is removed from the character matrix, *Homo* is inferred as monophyletic, *Au. garhi* as the sister taxon to *Paranthropus*, and *Au. africanus* as the sister taxon to the *Au. garhi*-*Paranthropus* clade. More nuanced approaches may include treating *K. platyops* as evidence of polymorphism in a variety of *Australopithecus* OTUs [45]. However, given that this taxon and its taphonomically-distorted morphology are somewhat contentious [45,46], until more and better preserved fossil material is recovered, any character matrix that includes *K. platyops* is inherently predicating their inferred topology on the taxonomic validity of *K. platyops*.

## Conclusions

We assessed the phylogenetic information content of 107 hominin craniodental characters and found that only 23 are uniquely phylogenetically informative. We document a disconnect in the dataset between a high cladistic information content and the direct assessment of what phylogenetic information a trait contributes to a topology inferred using the entire matrix. However, phylogenetic information as defined here is not a valuation of utility but an assessment of a trait’s contribution towards our specific baseline topology.

We also assessed the amount of information contained in skull anatomical units and found that only the maxilla contains significantly more information than expected. Nevertheless, basing systematic arguments solely on isolated units can lead to substantial inconsistencies in the inferred topologies: on its own, the maxilla and all other AUs are insufficient for accurate phylogenetic inference, at least based on character matrix and methods used here. While likelihood-based methods may allow more precise filtering of tree space, parsimony alone is insufficient to distinguish between potentially millions of most parsimonious phylogenetic trees when using single AUs.

Finally, we argue that the inclusion of certain taxa in phylogenetic analyses may strongly bias the inferred topology. Our concerns are consistent with the challenges of taxonomic attribution of new fossil specimens in paleoanthropology and suggestive that most fossil hominin taxa should be considered tentative [50]. Phylogenetic placements of taxa that are represented primarily by isolated anatomical units should be regarded as putative until the following prerequisites are met. First, phylogenetic inference accuracy may be improved by including more specimens (or more complete specimens) and OTUs. Second, we should incorporate more informative characters beyond those included in this dataset, including traits that are informative across the great apes rather than defined based on hominins alone, thus avoiding circularity. Lastly, we need to increase the application of likelihood-based methods to paleoanthropological systematics and have a better understanding of the information content of traits already in use, as we, and others [6] have done here.

Phylogenetic inference that aims to infer the most accurate species tree should only occur after having identified and removed problematic or uninformative traits and that systematic hypotheses be tempered not solely based on the confidence in the resolved tree (e.g., node support values), but also by the quality of the character matrix. The relationship between traits and phylogenies explored here ultimately carries implications across paleoanthropology, from taxonomic identification to the systematics of the entire hominin clade. Moreover, our methodology is flexible and potentially generalizable to other extinct or mostly-extinct clades beyond hominins.

## Materials and Methods

### Dataset and Phylogenetic Trees

The dataset used here [14] contains 107 discrete craniodental characters covering the entirety of the cranium across 20 OTUs, of which 13 are extinct and seven extant (SOM Table S2): *Colobus* (used to root the tree), *Papio, Hylobates, Pongo pygmaeus, Gorilla gorilla, Pan troglodytes, Sahelanthropus tchadensis, Ardipithecus ramidus, Au. anamensis, Au. afarensis, Au. africanus, Au. garhi, P. aethiopicus, P. robustus, P. boisei, Kenyanthropus platyops, Homo rudolfensis, H. habilis, H. ergaster*, and recent *H. sapiens*. We use the terms “trait” and “character” interchangeably.

All phylogenetic analyses were done in R v4.4.1 (Development Core Team R 2009) using the packages ape [51], TreeTools [52], and phangorn [53]. We inferred a baseline phylogenetic tree topology from Mongle et al. [14]‘s full dataset (SOM Table S1) as well as test tree topologies from subsets or modified versions of the character matrix. We used the branch and bound algorithm (BAB, [54]), which guarantees returning the MP tree(s), and maximum clade credibility as implemented in phangorn to construct the consensus trees. We inferred all topologies (except those where we were testing for the effects of each OTU) using all 20 OTUs and, while all trees are depicted in figures as rooted with *Colobus*, the ingroup topology is the same whether rooting with *Colobu*s or *Papio*. Topological distances were calculated on unrooted topologies and only on the hominin + *Pan troglodytes* subtree. All tree topologies were plotted with ggtree package [55].

### Phylogenetic information content and redundancy metrics

We used the cladistic information content [18] metric (*CIC*, as implemented in the TreeTools package) as a quantifier of whether a trait has intrinsic phylogenetic information content (PHIC), regardless of other characters and the character’s fit to any proposed tree. This metric calculates information by taking the negative natural logarithm of the proportion of binary trees (rooted trees with exactly two descendant lineages from each node) for which the trait of interest is homoplasy-free [18,56,57]. For a trait with either a unique or an identical state for every tip, this proportion is 1 and *CIC* = 0. In contrast, a trait with few states shared across several tips would have a relatively high *CIC*. Because *CIC* is a relative metric with no fixed upper bound, here we discuss a trait’s *CIC* as its rank (*CIC*_*rank*_) in relation to the value for all traits. We resolved ties by using the minimum position: if two traits have the fourth largest *CIC* value, they both have *CIC*_*rank*_ = 0. 97 and the fifth largest *CIC* value has *CIC*_*rank*_ = 0. 95. We also binned traits as defined “low *CIC*” (*CIC*_*rank*_ < 0. 25), “mid *CIC*” (0. 25 ≤ *CIC*_*rank*_ < 0. 75), and “high *CIC*” (0. 75 ≤ *CIC*_*rank*_).

We devised six complementary approaches (see Table 1) to assess the PHIC of effect of each character, anatomical unit (AU), and OTU on the baseline topology. Briefly, we either permuted individual trait values across OTUs, or removed traits and OTUs from the character matrix altogether. We then inferred the test tree from the modified character matrix, and computed the topological distance between each test topology and the baseline topology according to Robinson and Foulds [58] as implemented in the ape package.

### Permutation Robinson-Foulds distance (PRF)

For each of the 107 characters (one at a time), we permuted the character values across the hominin OTUs by sampling without replacement, restricting permutations solely to the hominin clade and measuring *RF* within the hominin +*Pan* subtree topology. Including extant outgroups in permutations *RF* calculations between test and baseline topologies might risk identifying traits that are informative on a higher taxonomic level (e.g., across catarrhines) and not necessarily within hominins. We then 1) inferred the MP tree(s) and the consensus topology from the modified character matrix, and 2) computed *RF* between the baseline and each of the test topologies. We repeated this process several times to generate a distribution. We compared results with 100, 250 and 500 permutations for five traits. The lowest *p*-value was for the comparison between 100 and 500 permutations (Mann-Whitney U, *p* = 0. 059, SOM Table S4), so we used 250 permutations. Permuting a character before inferring the tree topology allowed us to directly test how variability in that character affects the phylogenetic position of the tips. If a character was allowed to vary across the character matrix and those permutations created a distribution with many changes to the topology of the resulting tree, we concluded that the observed character distribution contains substantial information for the inferred clustering of tips. We refer to the distribution of these values as the PRF distribution (Table 1). For cross-trait comparisons, we used the median of this distribution 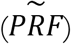 and the range between the 25% and 75% percentiles (interquartile range, *PRF*_*IQR*_) to characterize the spread of the PRF distribution.

### Removal RF (RRF)

We removed each of the 107 characters (one at a time) from the character matrix, inferred a test topology, and calculated calculated its RF distance to the baseline topology. This value (*RRF*) was compared across traits. We devised RRF to assess to what degree of uniqueness of the PHIC contributed by a character to the baseline topology. We tested for correlation between the different metrics (*CIC*, 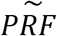, *RRF*) for each trait using Kendall’s rank correlation. Using both PRF and RRF, we classified traits in terms of their PHIC. We considered traits with 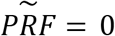 uninformative, traits with 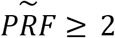 or *RRF* ≥ 2 uniquely informative, and traits with 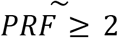 and *RRF* = 0 informative, but redundant. *RF* = 2 represents one change in the number of partitions created from the internal branches— i.e., the most minor non-polytomic topological change [58,59]. For example, the topology of *Paranthropus* in our baseline topology (Figure 1) has *P. aethiopicus* as the outgroup to *P. boisei* and *P. robustus*. If shuffling a trait’s values (and leaving all others intact) caused *P. boisei* to become the outgroup to *P. aethiopicus* and *P. robustus*, we would have *RF* = 2 and consider the trait as phylogenetically informative. But if that change resulted in a polytomy (*RF* = 1) or did not change the topology at all, we considered that trait as uninformative. Similarly, if the removal of a character did not affect the resulting tree (*RRF* = 0) even though its permutation did 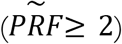, we inferred that the phylogenetic information present in that character is redundant, meaning the information was replicated in other traits and still present when said trait was removed from the inference.

### Unit removal Robinson-Foulds distance (URRF)

We carried out the same steps described for RRF but by removing all traits within an AU simultaneously. Traits were grouped based on location in the broader cranial structure into one of ten possible AUs (Table 2): frontal, nasal, maxilla, mandible, dentition, zygomatic, temporal, parietal, occipital, and “complex” (traits involving more than one bone). This approach allowed us to assess the impact on the inferred phylogeny of removing a set of traits more likely to not be independent.

### Single unit Robinson-Foulds distance” (SURF)

Here we instead used solely the traits in a given AU to infer a test topology. This approach allowed us to assess the relative weight of these AUs on the resulting hominin topology and to test whether the information contained in isolated fossil finds consisting of a single element is sufficient to resolve any recognizable clades from the baseline topology.

To test whether an AU had an unexpected amount of PHIC, we generated SURF and URRF null distributions by randomly sampling 100 random sets of *n* traits from the dataset, where *n* is the number of traits in a given AU (or 107 minus that number for URRF) and contrasted our empirical values with their respective distributions. We report two-tailed *p*-values and assumed α = 0. 05 for significance testing and, for significant cases, we also report the direction of the effect and Bonferroni adjusted *p*-values for multiple comparisons. We did not report SURF or URRF values for the nasal AU due to its consisting of a single character (trait 1, SOM Tables S1 and S5). We also used this approach to assess whether the *RF* between other test trees (the informative traits set and the uniquely informative set) and the baseline topology was unexpected.

### Taxon removal RF (TRRF)

Similar to RRF and used to measure the effect of the presence of an OTU on the inferred test tree topology. This approach required extra care to ensure compatibility when comparing our test tree (inferred with 19 OTUs) and baseline topology (20 OTUS). For *Colobus, Papio, Hylobates, Po. pygmaeus*, and *G. gorilla*, because we already limit all RF calculations to the 15 OTUs in the hominin + *Pan* subtree, we were able to directly compare the TRRF test topology (inferred with 19 OTUs, 15 included in the RF calculation) to the baseline. For hominins and *Pan*, we dropped those taxa from the baseline topology and then calculated *RF*. E.g., the TRRF test topology for *P. boisei* was inferred with 19 OTUS (14 used in calculating RF) and compared against the baseline tree but dropping *P. boisei* from the *Pan* + hominin subtree for the RF calculation.

## Supporting information

SOM_File1

SOM_table_s5

## Data Availability

All statistical analyses and figures were performed in R. Scripts necessary to reproduce the present results are available in the electronic supplementary material.

## Electronic Supplementary Material

SOM_File1.pdf (Figures S1-S23, Tables S1-S4, S6-S8)

SOM_File2.zip (code)

SOM_table_S5.xlsx

## Acknowledgments

We are grateful to Nicholas Post and Ward Wheeler for their helpful comments about heuristic *vs*. non-heuristic algorithms and to Theodore Schurr for his comments on an early version of this manuscript. We are especially appreciative of Carrie Mongle and collaborators for their commitment to sharing their 2019 dataset openly, and we thank Carrie Mongle for her helpful clarifications about the minutiae of the 2019 dataset used in our study.

## Notes

### Competing Interest Statement

The authors have declared no competing interest.

### Summary of Updates

This new version is revised to update the title and abstract, as well format for submission.

