## Supplementary material for "Assessing phylogenetic information content and redundancy in hominin craniodental traits": SOM_File1

### Supplementary Online Material

#### Supplementary Tables

**Table S1. Characters used in the analysis.** The following description of the characters and how each is coded in Mongle et al. (2019) serves as a reference for readers. Retrieved from Morphobank (http://morphobank.org/permalink/?P3151, accessed March 2023). In the main text, characters are referred to by their index number shown in the first column. Note: this table spans multiple pages.

| Character | Description | State |
|---|---|---|
| 1. SG1 | Projection of nasal bones above frontomaxillary suture | Project tapered (0); Project expanded (1); Not Projected (2); Variable (3) |
| 2. SG2 | Inferior orbital margin rounded | No (0); Variable (1); Yes (2) |
| 3. SG3 | Infraorbital foramen location | High (0); Variable (1); Low (2) |
| 4. SG4 | Anterior pillars | Absent (0); Variable (1); Present (2) |
| 5. SG5 | Nasoalveolar clivus contour in coronal plane | Convex (0); Straight (1); Concave gutter (2) |
| 6. SG6 | Protrusion of incisor alveoli beyond bicanine line basal view | Yes (0); No (1) |
| 7. SG7 | Nasal cavity entrance | Stepped overlap (0); Smooth overlap (1); Smooth no overlap (2); Stepped no overlap (3); Variable stepped with or without overlap (4) |
| 8. SG8 | Palate thickness | Thin (0); Thick (1) |
| 9. SG9 | Height of the masseter origin | Low (0); Variable (1); High (2) |
| 10. SG10 | M-L (mediolateral) thickness of zygomatic arch at root of frontal process | Thin (0); Thick (1) |
| 11. SG11 | Anterior projection of zygomatic bone relative to piriform aperture dishing | Posterior (0); Variable Posterior Intermediate (1); Intermediate (2); Variable Intermediate Anterior (3); Anterior dished (4) |
| 12. SG12 | Anterior palatal depth | Shallow (0); Variable (1); Deep (2) |
| 13. SG13 | Index of palate protrusion anterior to sellion facial prognathism | Hyper prognathic (0); Prognathic (1); Variable prog |
| 14. SG14 | Masseteric position relative to sellion | At or posterior (0); Variable (1); At or anterior (2) |
| 15. SG15 | Maxillary trigon zygomaticomaxillary step | Absent (0); Variable (1); Present (2) |
| 16. SG16 | Cranial capacity | Small (0); Intermediate (1); Large (2); Very Large (3) |
| 17. SG17 | Cerebellar morphology | Lateral flare posterior protrusion (0); Tucked (1) |

| | | |
|---|---|---|
| 18. SG18 | O-M[1] sinus present in high frequency | No (0); Intermediate (1); Yes (2) |
| 19. SG19 | Anteromedial incursion of the superior temporal lines | Weak (0); Variable from Moderate to Weak (1); Moderate (2); Strong (3) |
| 20. SG20 | Sagittal crest present at least in presumptive males | Yes (0); No (1) |
| 21. SG21 | Compound T-N (temporonuchal) crest at least in presumptive males | Extensive (0); Variable (1); Partial (2); Absent (3) |
| 22. SG22 | Asterionic notch | Present (0); Variable (1); Absent (2) |
| 23. SG23 | Parietal overlap of occipital at asterion at least in males | No (0); Yes (1) |
| 24. SG24 | Squamosal suture overlap extensive at least in males | Not extensive (0); Extensive (1) |
| 25. SG25 | Lateral inflation of mastoid process relative to supramastoid crest | Not inflated (0); Variable (1); Inflated (2) |
| 26. SG26 | Postorbital constriction | Marked (0); Moderate (1); Slight (2) |
| 27. SG27 | Pneumatization of temporal squama | Extensive (0); Variable (1); Reduced (2) |
| 28. SG28 | Facial hafting[2] | Low (0); High (1) |
| 29. SG29 | Supraglenoid gutter width | Narrow (0); Wide (1) |
| 30. SG30 | External cranial base flexion | Retroflexed (0); Flat (1); Moderate (2); Flexed (3) |
| 31. SG31 | Horizontal distance between temporomandibular joint (TMJ) and upper M2/M3 | Long (0); Short (1) |
| 32. SG32 | Relative depth of mandibular fossa | Shallow (0); Variable between shallow and intermediate (1); Intermediate (2); Deep (3) |
| 33. SG33 | Postglenoid process size | Large (0); Mid sized (1); Variable mid |
| 34. SG34 | Configuration of tympanic | Tubular or weak crest (0); Crest with vertical plate (1); Crest with inclined plate (2) |
| 35. SG35 | Mediolateral position of external auditory meatus | Medial (0); Variable (1); Lateral (2) |
| 36. SG36 | Vaginal process[3] | Small or absent (0); Variable (1); Moderate to large (2) |
| 37. SG37 | Eustachian process of tympanic | Present and prominent (0); Variable (1); Absent or slight (2) |
| 38. SG38 | Petrous orientation | Sagittal (0); Intermediate (1); Coronal (2) |
| 39. SG39 | Heart-shaped foramen magnum | Absent (0); Variable (1); Present (2) |

[1] Occipitomarginal sinus, cf. Strait et al., 1997 (Mongle, pers. comm.)

[2] cf. Strait et al., 1997 (Mongle, pers. comm.)

[3] of the tympanic, cf. Strait et al., 1997 (Mongle, pers. comm.)

| | | |
|---|---|---|
| 40. SG40 | Inclination nuchal plane | Extremely steep (0); Steeply inclined (1); Intermediate (2); Variable (3); Weakly inclined (4) |
| 41. SG41 | Position of foramen magnum relative to bi-tympanic line | Well posterior (0); At bi tympanic line (1); Variable at or anterior (2); Well anterior (3) |
| 42. SG42 | Inclination of foramen magnum | Strongly inclined posterior (0); Roughly horizontal (1); Strongly inclined anterior (2) |
| 43. SG43 | Origin of digastric muscle | Broad shallow fossa (0); Deep narrow notch (1) |
| 44. SG44 | Mandibular cross sectional area at M1 | Small (0); Variable (1); Large (2) |
| 45. SG45 | Orientation of mandibular symphysis | Receding (0); Intermediate Variable (1); Vertical (2) |
| 46. SG46 | Direction of mental foramen opening | Anterior (0); Variable (1); Lateral (2); Posterior (3) |
| 47. SG47 | Hollowing above and behind mental foramen | Present (0); Variable (1); Absent (2) |
| 48. SG48 | Width of mandibular extramolar sulcus | Narrow (0); Variable (1); Wide (2) |
| 49. SG49 | Mandibular deciduous canine shape | Apex central, mesial convexity low (0); Apex mesial, mesial convexity high (1) |
| 50. SG50 | Incisal reduction | No (0); Moderate (1); Yes (2) |
| 51. SG51 | Canines reduced | No (0); Somewhat (1); Very (2) |
| 52. SG52 | Prominence of median lingual ridge of mandibular canine | Prominent (0); Variable (1); Weak (2) |
| 53. SG53 | Premolar crown area | Smallest (0); State 1 (1); State 2 (2); State 3 (3); State 4 (4); Largest (5) |
| 54. SG54 | Molar crown area | Smallest (0); State 1 (1); State 2 (2); Largest (3) |
| 55. SG55 | Lower dm1 mesial crown profile | Mesial marginal ridge (MMR) absent, protoconid anterior fovea open (0); MMR slight, protoconid anterior fovea open (1); MMR slight, protoconid anterior fovea closed (2); MMR thick, protoconid even with metaconid fovea closed (3) |
| 56. SG56 | Distal marginal ridge of upper dm2 | Low (0); High (1) |
| 57. SG57 | Positions of buccal and lingual cusps relative to crown margin (states as for mandibular teeth, reverse for maxillary teeth) | Buccal and lingual cusps approximate crown margin (0); Lingual cusps approximate margin, buccal cusps slightly lingual to margin (1); Lingual cusps approximate margin, buccal cusps moderately lingual to margin (2); Lingual cusps slightly buccal to margin, buccal cusps moderately lingual to margin (3); Lingual cusps moderately buccal to margin, buccal cusps strongly lingual to margin (4) |
| 58. SG58 | Frequency of well developed lower P3 metaconid | Absent (0); Infrequent (1); Frequent (2) |
| 49. SG59 | Relative enamel thickness | Thin (0); Variable (1); Thick (2); Hyperthick (3) |
| 60. SG60 | Dental development rate | Delayed (0); Intermediate (1); Accelerated (2) |

| | | |
|---|---|---|
| 61. SG61 | Mesiobuccal protrusion of P3 crown base | Strong (0); Moderate (1); Variable (2); Weak to absent (3) |
| 62. SG62 (=CW86) | Orientation of mandibular premolar row dental arcade shape | Parasagittal (0); Oblique (1) |
| 63. SG63 | Parietal tuber | Absent (0); Present (1) |
| 64. SG64 | Parietomastoid angle | Strong (0); Weak (1) |
| 65. SG65 | External auditory meatus size | Small (0); Large (1) |
| 66. SG66 | Separation of mandibular tooth rows | Widely Separated (0); Narrow Separation (1) |
| 67. SG67 | Configuration of the superior orbital fissure | Foramen (0); Comma shaped (1) |
| 68. SG68 | Size of Longus capitis insertion | Large (0); Small (1) |
| 69. SG69 | Height of articular eminence above occlusal plane | High above plane (0); Near the plane (1) |
| 70. SG70 (=CW38) | Extensive mesial groove on upper canine | Yes (0); No (1) |
| 71. MSG1 | Supraorbital hollowing | Absent (0); Variable Absent to restricted midline (1); Restricted midline delimited depression (2); Restricted lateral depression (3); Supratoral sulcus that extends from midline laterally beyond medial third of orbit (4) |
| 72. CW1 | Depth of subarcuate fossa | Deep (0); Moderately deep to shallow (1); Very shallow to absent (2) |
| 73. CW2 | Orientation and length of the post incisive planum | Long weakly inclined (0); Intermediate (1); Short steeply inclined (2) |
| 74. CW3 | Distinctiveness of angular process of mandible | Distinct with posterior projection (0); Not Distinct (1) |
| 75. CW17 | Presence/absence of frontal sinus | Absent (0); Present (1) |
| 76. CW19 | Position of infraorbital foramen relative to orbit | Foramen beneath middle third of orbital breadth (0); Foramen beneath medial third of orbital breadth (1) |
| 77. CW20 | Orientation of zygomatic bone | Frontal (0); Variable (1); Fronto lateral (2); Lateral (3) |
| 78. CW22 | Glabellar prominence relative to sella | Strong, projects anterior to sella (0); Intermediate, at the level of sella (1); Weak, recessed behind sella (2) |
| 79. CW25 | Supraorbital expression | Weak (0); Intermediate (1); Torus-like (2) |
| 80. CW26 | Supraorbital contour | Arched (0); Variable (1); Less arched (2) |
| 81. CW32 | Position of zygomatic foramina | At or below plane of orbital rim (0); Above plane of orbital rim (1) |
| 82. CW35 | Patency of premaxillary suture in adults from frontal view | Patent (0); Variable (1); Obliterated (2) |

| | | |
|---|---|---|
| 83. CW36 | Petrous apex ossified beyond sphenooccipital synchondrosis | Not ossified (0); Ossified with projection (1) |
| 84. CW40 | Upper I2 similar in shape to upper I1 | Dissimilar (0); Variable (1); Similar (2) |
| 85. CW41 | Robusticity of canines | Slender (0); More robust (1) |
| 86. CW42 | Basal keel of lower canine | Present (0); Reduced (1); Absent (2) |
| 87. CW43 | Basal area of paracone of P3 | Paracone much larger than protocone (0); Paracone larger than protocone (1); Paracone equals protocone (2) |
| 88. CW46 | Metaconid of lower dp3 | Absent or poorly defined (0); Well defined (1) |
| 89. CW48 | Talonid basin of lower dp3 | Open distally (0); Closed distally (1) |
| 90. CW50 | Distal trigonid crest on lower dp4 | Does not reach protoconid apex (0); Reaches protoconid apex (1) |
| 91. CW52 | Protocone of upper dp3 in occlusal view | Larger than paracone (0); Smaller than paracone (1) |
| 92. CW54 | Crista obliqua of upper dp4 | Weak (0); Moderate (1); Strong (2) |
| 93. CW59 | Insertion of genioglossus | Above inferior transverse torus (0); On inferior transverse torus (1) |
| 94. CW60 | Insertion[4] of geniohyoideus | Basally on inferior transverse torus (0); Higher on inferior transverse torus (1); Above inferior transverse torus (2) |
| 95. CW61 | Insertion[5] of digastric | Posterior to inferior transverse torus (0); Inferior transverse torus (1); Variable (2); Not on symphysis (3) |
| 96. CW64 | Condylar canal | Absent or infrequent (0); Frequently present (1) |
| 97. CW76 | Fovea posterior[6] | Absent or weak (0); Well developed (1) |
| 98. CW78 | Mandibular corpus depth along tooth row | Shallow mesially (0); Constant (1); Variable (2); Deepens mesially (3) |
| 99. CW89 | Upper I1 lingual crenulations | Absent (0); Marginal (1); Whole Surface (2) |
| 100. KRJ6 | Relative height of vertex | Low (0); Variable (1); Intermediate (2); High (3) |
| 101. KRJ11 | Midfacial prognathism | High (0); Intermediate (1); Low (2) |
| 102. KRJ13 | Position zygomatic angle | Below orbit (0); At or below (1); At orbit (2); At or above (3); Above (4) |
| 103. KRJ25 | Interorbital width | Broad (0); narrow (1) |
| 104. KRJ34 | Anterior pole shape[7] | Rounded (0); beaked (1) |

[4] In actuality, the origin of the genioglossus muscle, as it is assessed on the mandible (Mongle, pers. comm.)
[5] In actuality, the origin of the digastric muscle, as it is assessed on the mandible (Mongle, pers. comm.)
[6] Refers to the post talonid basin of the lower molars, cf. Strait & Grine, 2004, (Mongle, pers. comm.)
[7] Cf. Dean et al., 2000 (Pan and Gorilla only), Kimbel et al., 2004 (Mongle, pers. comm.)

| | | |
|---|---|---|
| 105. KRJ36 | Supraorbital corner | Rounded (0); Variable (1); Squared with weak or absent tubercle (2); Squared with prominent tubercle (3) |
| 106. KRJ58 | Nuchal plane form | Transversely convex (0); variable (1); flat (2) |
| 107. KRJ66 | Zygomatic frontal process lateral margin | Vertical (0); Slightly divergent (1); Strongly divergent (2) |

**Table S2. Taxonomic units from the Mongle et al. (2019) dataset and a summary of the degree of missing data for each taxon.** Taxa are ordered from earliest hominins to extant taxa, and from least to most missing data within each genus. The number of missing traits are listed per anatomical unit. Trait indexes correspond to those in SOM Table S1. See also: SOM Table S3 and Figure 1 (main text). Note: this table spans multiple pages.

| OTU | Number of Missing Traits | Missing traits |
|---|---|---|
| *Sahelanthropus tchadensis* | $n = 58$ | 58 traits: frontal ($n = 1$), nasal ($n = 1$), maxilla ($n = 6$), mandible ($n = 9$), dental ($n = 16$), zygomatic ($n = 3$), temporal ($n = 8$), parietal ($n = 5$), occipital ($n = 4$), complex ($n = 4$) |
| *Ardipithecus ramidus* | $n = 24$ | 24 traits: frontal ($n = 4$) maxilla ($n = 3$), mandible ($n = 1$), dental ($n = 5$), temporal ($n = 1$), parietal ($n = 3$), occipital ($n = 5$), complex ($n = 2$) |
| *Australopithecus africanus* | $n = 0$ | — |
| *Au. afarensis* | $n = 4$ | frontal ($n = 1$), zygomatic ($n = 1$), temporal ($n = 1$), occipital ($n = 1$) |
| *Au. anamensis* | $n = 75$ | 75 characters traits: frontal ($n = 9$), nasal ($n = 1$), maxilla ($n = 11$), mandible ($n = 4$), dental ($n = 10$), zygomatic ($n = 7$), temporal ($n = 10$), parietal ($n = 5$), occipital ($n = 9$), complex traits ($n = 9$) |
| *Au. garhi* | $n = 78$ | 78 traits: frontal ($n = 3$), nasal ($n = 1$) maxilla ($n = 5$), mandible ($n = 15$), dental ($n = 15$), zygomatic ($n = 6$), temporal ($n = 14$), parietal ($n = 3$), occipital ($n = 9$), complex ($n = 7$) |
| *Kenyanthropus platyops* | $n = 79$ | 79 traits: frontal ($n = 6$), nasal ($n = 1$), maxilla ($n = 5$), mandible ($n = 14$), dental ($n = 22$), zygomatic ($n = 5$), temporal ($n = 11$), parietal ($n = 4$), occipital ($n = 6$), complex ($n = 5$) |
| *Paranthropus boisei* | $n = 2$ | 2 dental traits |
| *P. robustus* | $n = 5$ | 5 traits: frontal unit ($n = 1$), mandible ($n = 1$), temporal ($n = 2$), occipital ($n = 1$) |
| *P. aethiopicus* | $n = 27$ | 27 traits: frontal ($n = 2$), maxilla ($n = 2$), mandible ($n = 2$), dental ($n = 15$), zygomatic ($n = 2$), temporal ($n = 1$), parietal ($n = 1$), occipital ($n = 2$) |
| *Homo sapiens* | $n = 0$ | — |
| *H. habilis* | $n = 6$ | 6 traits: mandible ($n = 3$), dental ($n = 3$) |

| | | |
|---|---|---|
| *H. ergaster* | $n = 9$ | 9 traits: frontal ($n = 2$), dental ($n = 4$), temporal ($n = 2$), occipital ($n = 1$) |
| *H. rudolfensis* | $n = 30$ | 30 traits: frontal ($n = 2$), maxilla ($n = 2$), mandible ($n = 1$), dental ($n = 9$), zygomatic ($n = 2$), temporal ($n = 7$), occipital ($n = 5$), complex ($n = 2$) |
| *Pan troglodytes* | $n = 0$ | — |
| *Gorilla gorilla*[a] | $n = 0$ | — |
| *Pongo pygmaeus*[a] | $n = 0$ | — |
| *Hylobates*[a] | $n = 1$ | 1 occipital traits |
| *Colobus* | $n = 2$ | 2 traits: dental ($n = 1$), occipital ($n = 1$) |
| *Papio* | $n = 2$ | 2 traits: dental ($n = 1$), occipital ($n = 1$) |

**Table S3. Number of OTUs coded for each character in the Mongle et al. (2019) dataset.** In parentheses are the total number of characters per number of OTUs coded. Trait indexes correspond to those in SOM Table S1. See also: SOM Table S3 and Figure 1 (main text).

| Number of OTUs coded | Traits |
|---|---|
| $n = 20$ | 51, 54, and 59 ($n = 3$) |
| $n = 19$ | 4, 8, 11, 13, 16, 19, 26, 32, 50, 53, and 57 ($n = 11$) |
| $n = 18$ | 2, 5, 6, 12, 15, 27, 28, 34, 35, 44, 46, 48, 61, 62, 66, 79, 85, 87, and 103 ($n = 19$) |
| $n = 17$ | 7, 9, 14, 20, 21, 31, 33, 38, 45, 47, 58, 65, 71, 73, 78, 80, 101, 102, and 107 ($n = 19$) |
| $n = 16$ | 1, 3, 17, 25, 29, 37, 39, 40, 41, 55, 63, 64, 69, 70, 86, 95, 97, 98, 105, and 106 ($n = 20$) |
| $n = 15$ | 10, 22, 23, 24, 36, 42, 52, 76, 77, 88, 93, 94, 100, and 104 ($n = 14$) |
| $n = 14$ | 30, 43, 68, 74, 81, 82, 84, 96, and 99 ($n = 9$) |
| $n = 13$ | 56, 72, 75, 89, 90, and 92 ($n = 6$) |
| $n = 12$ | 18, 49, 60, 67 ($n = 4$) |
| $n = 11$ | 83 and 91 ($n = 2$) |

**Table S4. Comparison of Robinson Foulds distance distributions for different numbers of permutations for five traits.** Trait indexes correspond to those in SOM Table S1. Columns 2-4 list $\boldsymbol{p}$-values for the Mann-Whitney U test.

| Trait | 100 *vs.* 250 permutations | 100 *vs.* 500 permutations | 250 *vs.* 500 permutations |
|---|---|---|---|
| 1 | 0.880 | 0.760 | 0.831 |
| 11 | 0.797 | 0.713 | 0.362 |
| 21 | 0.781 | 0.562 | 0.688 |
| 31 | 0.721 | 0.300 | 0.421 |
| 41 | 0.128 | 0.059 | 0.600 |

**Table S5. All statistics computed for each individual traits.**

See: SOM_table_S5.xlsx

**Table S6. Traits recommended for removal**. We include here 22 of the 76 uninformative traits ($\widetilde{PRF} = 0$) for which we never sampled a permuted test tree topology topologically different from the baseline topology (i.e., all 250 permuted test topologies were $RF = 0$ from the baseline). Trait indexes in parentheses correspond to those in SOM Table S1. *Character 83 has $CIC = 0$, the lowest observed in the dataset, thus its rank is 1/107.

| Trait | AU | $CIC_{rank}$ |
|---|---|---|
| Inclination of nuchal plane (40) | Occipital | 0.355 |
| Presence of an extensive mesial groove on $C^1$ (70) | Dentition | 0.533 |
| Depth of subarcuate fossa (72) | Temporal | 0.019 |
| Orientation and length of post incisive planum (73) | Mandible | 0.308 |
| Distinctiveness of angular process of mandible ( 74) | Mandible | 0.037 |
| Presence of frontal sinus (75) | Frontal | 0.243 |
| Position of infraorbital foramen relative to orbit (76) | Maxilla | 0.047 |
| Glabella prominence relative to sella (78) | Frontal | 0.280 |
| Position of zygomatic foramina (81) | Zygomatic | 0.150 |
| Patency of premaxillary suture in adults from frontal view (82) | Maxilla | 0.579 |
| Petrous apex ossified beyond sphenooccipital synchondrosis (83) | Temporal | 0.009* |
| Robusticity of canines (85) | Dentition | 0.393 |
| Basal area of paracone of P3 (87) | Dentition | 0.710 |
| Definition of metaconid of $dP_3$ (88) | Dentition | 0.495 |
| Open or closed talonid basin of $dP_3$ (89) | Dentition | 0.243 |
| Reach of distal trigonid crest on $dP_4$ (90) | Dentition | 0.299 |
| Size of protocone of $dP^3$ in occlusal view (91) | Dentition | 0.121 |
| Strength of crista obliqua of $dP^4$ (92) | Dentition | 0.215 |
| Location of the insertion of genioglossus (93) | Mandible | 0.159 |
| Location of insertion of the digastric (95) | Mandible | 0.654 |

| | | |
|---|---|---|
| Presence of fovea posterior (97) | Dentition | 0.308 |
| Presence of $I^1$ crenulations (99) | Dentition | 0.421 |

**Table S7. Uniquely informative traits**. Set of 23 traits that have $\boldsymbol{\widetilde{PRF} \geq 2}$ and $\boldsymbol{RRF \geq 2}$. Traits are sorted in descending order based on $\boldsymbol{\widetilde{PRF}}$ values. For these traits, $\boldsymbol{\widetilde{PRF}}$ and $\boldsymbol{RRF}$ are perfectly correlated, so the latter is omitted. Trait indexes in parentheses correspond to those in SOM Table S1.

| Trait | AU | $CIC_{ran}$ | $\widetilde{PRF}$ |
|---|---|---|---|
| Size of the postglenoid process (33) | Temporal | 0.963 | 10 |
| Prominence of the lingual ridge of the mandibular canine (52) | Dentition | 0.430 | 10 |
| Orientation of mandibular premolar row arcade shape (62) | Mandible | 0.701 | 10 |
| Flexion of external cranial base (30) | Occipital | 0.570 | 8 |
| Configuration of the tympanic (34) | Temporal | 0.589 | 8 |
| Orientation of the petrous (38) | Temporal | 0.888 | 8 |
| Midfacial prognathism (101) | Maxilla | 0.897 | 8 |
| Contour of the nasoalveolar clivus (5) | Maxilla | 0.869 | 6 |
| Protrusion of incisor alveoli beyond bicanine line (6) | Maxilla | 0.505 | 4 |
| Shape of nasal cavity entrance (7) | Maxilla | 0.794 | 4 |
| Anterior palatal depth (12) | Maxilla | 0.935 | 4 |
| Anteromedial incursion of the superior temporal lines (19) | Frontal | 0.869 | 4 |
| Horizontal distance between the temporomandibular joint and $M^2/M^3$ (31) | Complex | 0.206 | 4 |
| Orientation of the mandibular symphysis (45) | Mandible | 0.850 | 4 |
| Hollowing above and behind mental foramen (47) | Mandible | 0.776 | 4 |
| Supraorbital hollowing (71) | Frontal | 0.748 | 4 |
| Supraorbital expression (79) | Frontal | 0.738 | 4 |
| Anterior projection of zygomatic bone (11) | Zygomatic | 0.280 | 2 |
| Cerebellar morphology (17) | Complex | 0.533 | 2 |
| Pneumatization of temporal squama (27) | Temporal | 0.645 | 2 |
| Mesiobuccal protrusion of P3 crown base ( 61) | Dentition | 0.776 | 2 |

| | | | |
|---|---|---|---|
| Configuration of superior orbital fissure (67) | Frontal | 0.131 | 2 |
| Supraorbital corner (105) | Frontal | 0.804 | 2 |

**Table S8. Contribution of each OTU to the inferred topology**. TRRF denotes the Robinson-Foulds distance between the baseline tree and each test tree. For each OTU tested, the baseline tree was inferred using all taxa followed by dropping the tested OTU, and the test tree was inferred using a modified character matrix where the OTU was entirely removed. TRRF test trees for all taxa with $\boldsymbol{TRRF > 0}$ are shown in SOM Figures S9-S23, except *K. platyops,* shown in Figure 5 (main text).

| OTU | $TRRF$ |
|---|---|
| *Pan troglodytes* | 10 |
| *Au. anamensis* | 8 |
| *K. platyops* | 8 |
| *H. sapiens* | 8 |
| *H. rudolfensis* | 8 |
| *P. robustus* | 8 |
| *Au. afarensis* | 6 |
| *Au. africanus* | 6 |
| *H. habilis* | 6 |
| *Colobus* | 4 |
| *G. gorilla* | 4 |
| *Hylobates* | 4 |
| *H. ergaster* | 4 |
| *Papio* | 4 |
| *P. boisei* | 4 |
| *S. tchadensis* | 4 |
| *Ar. ramidus* | 0 |
| *Au. garhi* | 0 |
| *P. aethiopicus* | 0 |

| | |
|---|---|
| *Po. pygmaeus* | 0 |

#### Supplementary Figures

**Figure S1. Comparison between phylogenetic information content metrics using two different baseline trees.** In both figures, the x-axis shows Robinson-Foulds distances based on the baseline tree (Figure 1, main text) and the parsimony-derived topology (y-axis) from Mongle et al. (2019). A) Removal Robinson-Foulds distances ($\boldsymbol{RRF}$). B) Median permutation Robinson-Foulds distances ($\widetilde{\boldsymbol{PRF}}$). Both correlations are significant ($\boldsymbol{p < 0.001}$, Kendall's τ correlation).

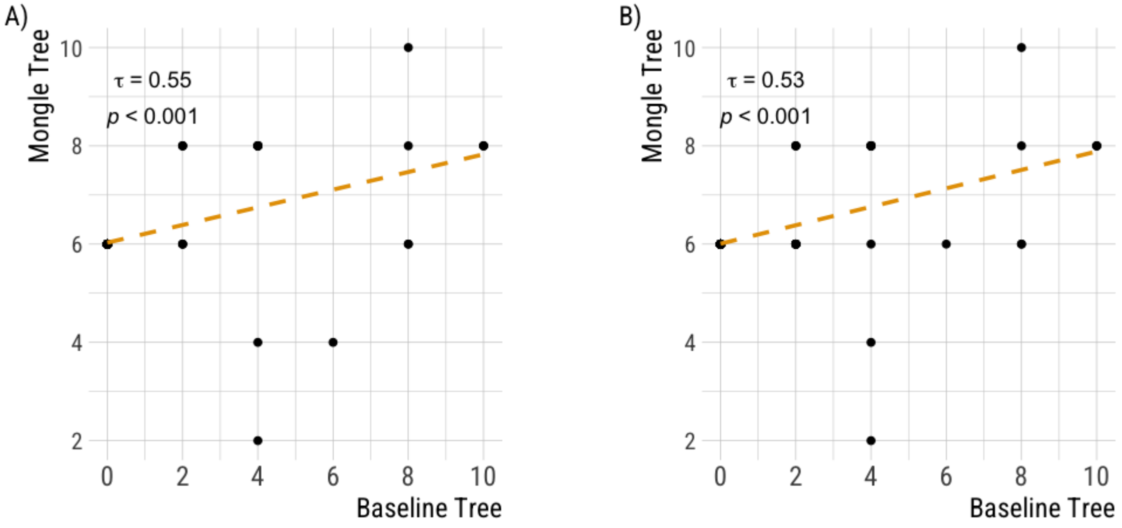

**Figure S2. Number of informative and uninformative traits in each anatomical unit (AU).** Raw two-tailed $\boldsymbol{p}$-values were obtained by comparing the proportion of uninformative to informative traits to the same proportion from 10,000 randomly drawn random sets of traits (pseudo-AUs) of the same length as each AU. All $\boldsymbol{p}$-adjusted (Bonferroni) values are $\geq \mathbf{0.135}$. The nasal unit ($\boldsymbol{n} = \mathbf{1}$) was omitted.

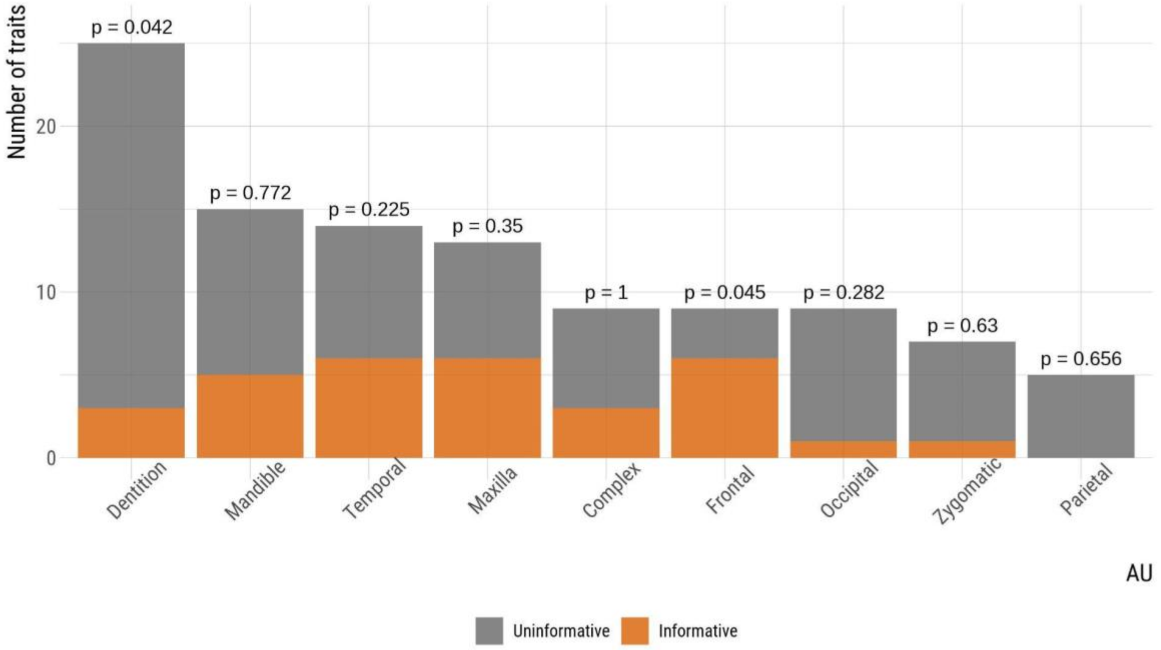

**Figure S3. Effect of character 33 (the size of the postglenoid process) on the baseline tree when removed or permuted.** Left, consensus tree topology inferred using the branch and bound algorithm with maximum clade credibility when character 33 (the size of the postglenoid process, $\boldsymbol{RRF = 10}$, $\boldsymbol{CIC_{rank} = 0.963}$) is removed from the character matrix. Right, the most commonly inferred consensus tree topology for the permutation of character 33 across the hominin OTUs ($\boldsymbol{RF = 10}$ from baseline tree for the permutation shown and $\widetilde{\boldsymbol{PRF}} \boldsymbol{= 10}$ across all permutations). Purple diamonds indicate all changes in relation to the baseline topology, but only the differences in the hominin + *Pan* subtree were considered in the RF calculation.

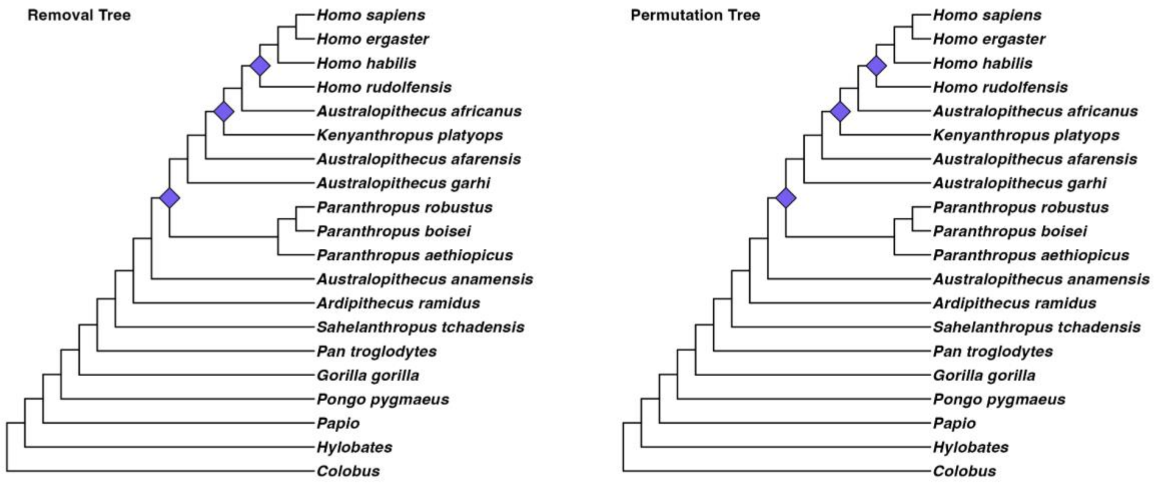

**Figure S4. Effect of character 52 (prominence of the lingual ridge of the $C_1$) on the baseline tree whe removed or permuted.** Left, consensus tree topology inferred using the branch and bound algorithm with maximum clade credibility when character 52 (prominence of the lingual ridge of the $C_1$, $\boldsymbol{RRF = 10}$, $\boldsymbol{CIC_{rank} = 0.430}$) is removed from the character matrix. Right, the most commonly inferred consensus tree topology for one permutation of character 52 across the hominin OTUs ($\boldsymbol{RF = 10}$ from baseline tree for the permutation shown and $\widehat{\boldsymbol{PRF}} = \boldsymbol{0}$ across all permutations). Purple diamonds indicate all changes in relation to the baseline topology, but only the differences in the hominin + *Pan* subtree were considered in the RF calculation.

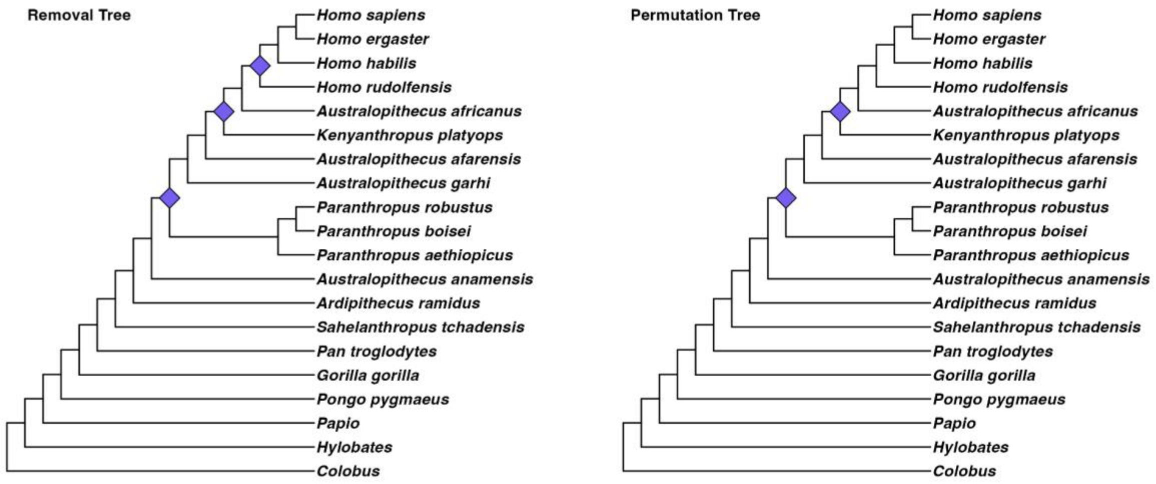

**Figure S5. Effect of character 62 (orientation of mandibular premolar row arcade shape) on the baseline tree whe removed or permuted. .** Left, consensus tree topology inferred using the branch and bound algorithm with maximum clade credibility when character 62 (orientation of mandibular premolar row arcade shape, $\boldsymbol{RRF = 10}$, $\boldsymbol{CIC_{rank} = 0.701}$) is removed from the character matrix. Right, the most commonly inferred consensus tree topology for one permutation of character 62 across  the hominin OTUs ($\boldsymbol{RF = 10}$ from baseline tree for the permutation shown and $\boldsymbol{\widetilde{PRF} = 0}$ across all permutations). Purple diamonds indicate all changes in relation to the baseline topology, but only the differences in the hominin + *Pan* subtree were considered in the RF calculation.

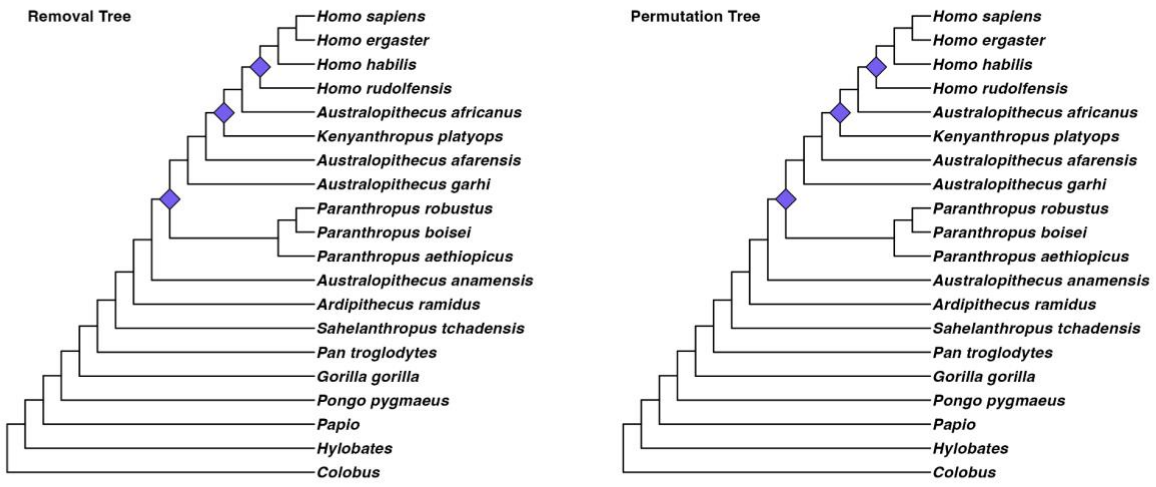

**Figure S6. Pairwise Robinson-Foulds distances between removal trees for 23 uniquely informative traits.** The matrix shows RF distances between all pairs of test trees inferred by removing each of the 23 uniquely informative traits from the character matrix. All RF distances are even (i.e., not polytomies), and the scale at the bottom of the figure shows their observed values. Trait indexes correspond to those in SOM Table S1.

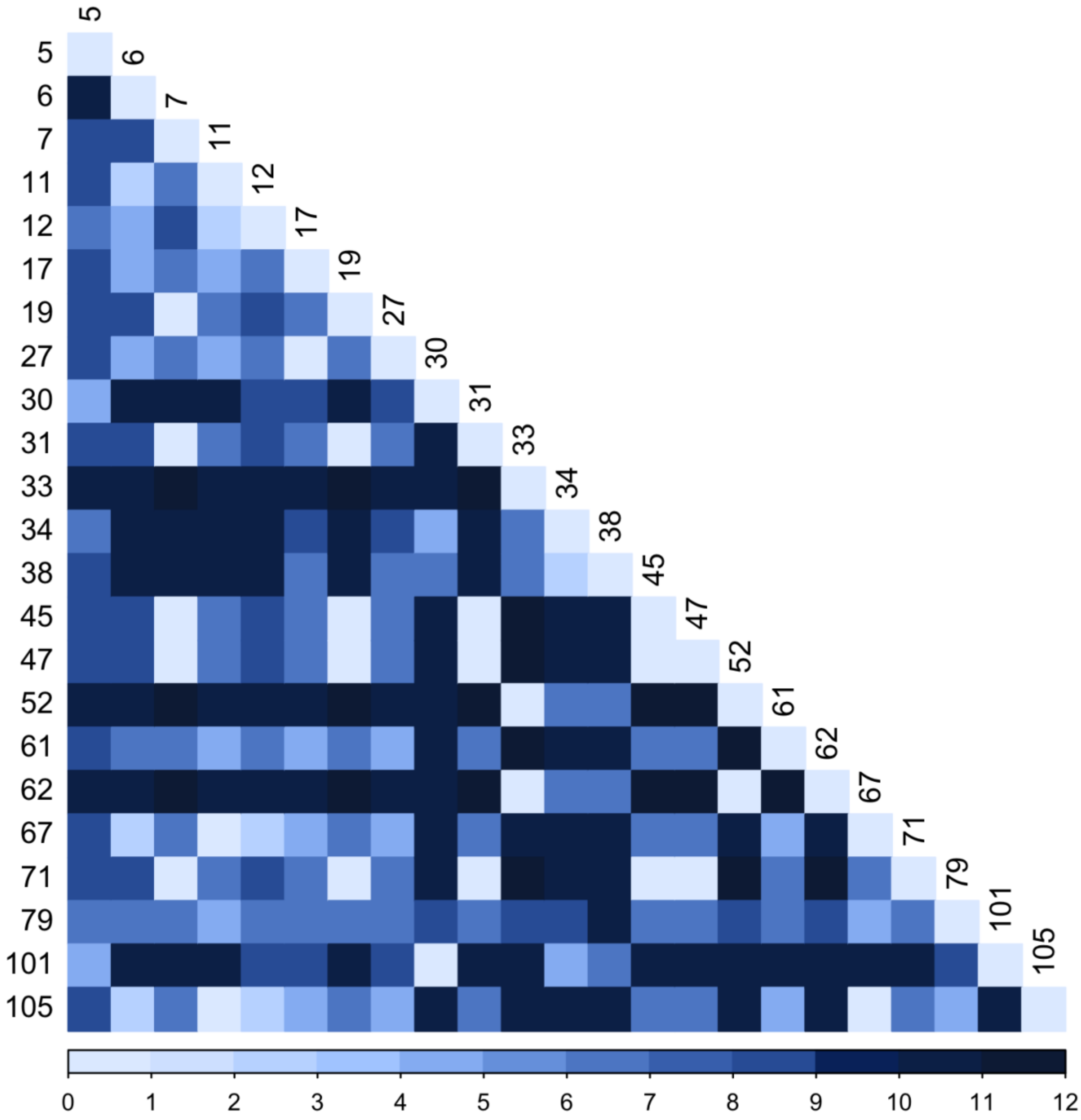

**Figure S7. Hominin phylogenetic tree inferred from 23 uniquely informative traits**. A) The consensus tree topology inferred from solely the 23 uniquely informative traits using the branch and bound algorithm with maximum clade credibility ($\boldsymbol{RF = 4}$ between this and the baseline topology). Purple diamonds indicate all changes in relation to the baseline topology, but only the differences in the hominin + *Pan* subtree were considered in the RF calculation. B) Null distribution of Robinson-Foulds distances between 100 random sets of 23 traits and the baseline tree ($\boldsymbol{p = 0.02}$; Figure 1, main text). The dashed orange line shows the empirical RF distance between the tree in A) and the baseline tree.

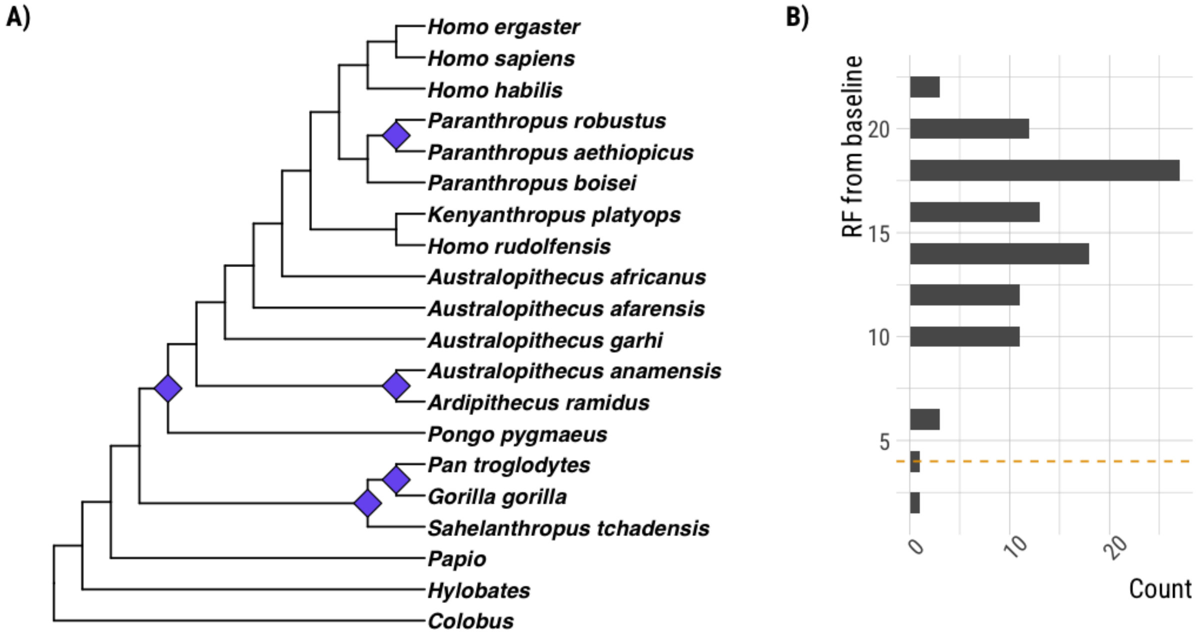

**Figure S8. Robinson Foulds distances and null distributions for each anatomical unit (AU)**. All figures show null distributions of Robinson-Foulds distances between test trees inferred from 100 random sets of $n$ traits and the baseline tree, where $n$ is either A) the number of traits in each AU or B) 107 minus the number of traits in each AU. The dashed orange lines show the empirical value observed for A) SURF and B) URRF. The "complex" AU includes traits that encompass more than one bone. The nasal AU is omitted due to only containing one trait (see Methods in main text). For both the complex and parietal units, the empirical SURF result could not be completed due to computational constraints. See also: Table 2 in main text.

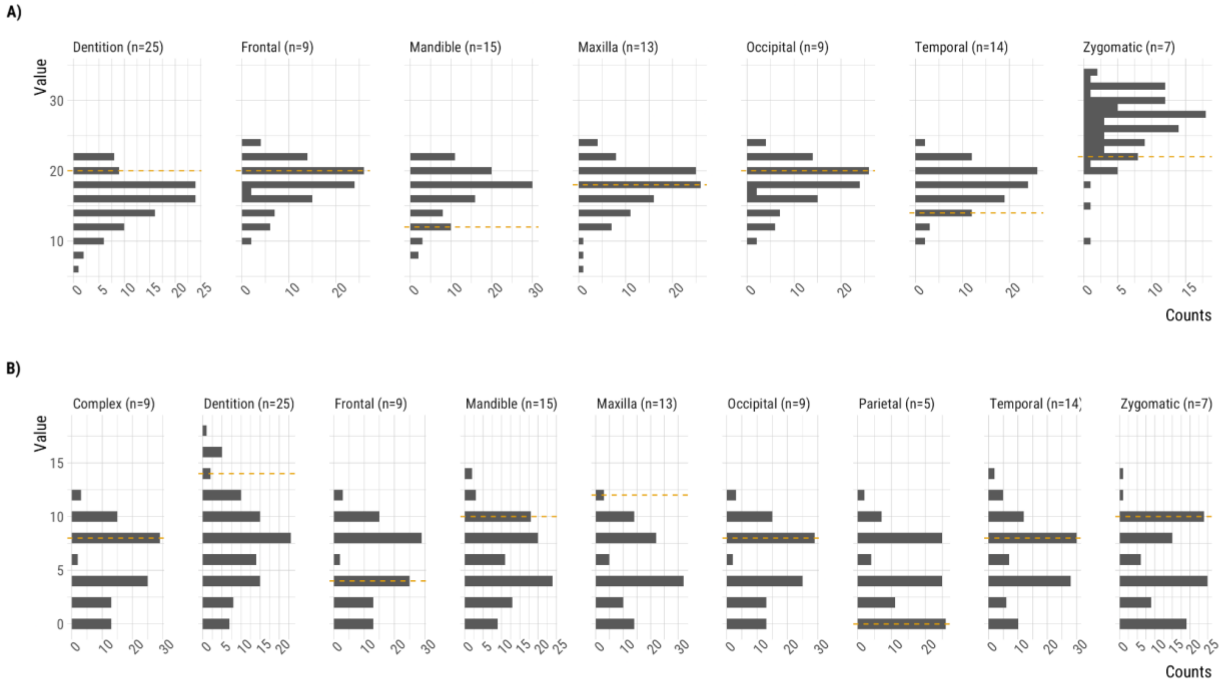

**Figure S9. Hominin phylogenetic tree topology inferred without *Pan troglodytes*.** The consensus tree topology inferred from the character matrix excluding *Pan troglodytes* using the branch and bound algorithm with maximum clade credibility ($\boldsymbol{RF = 10}$ between this and the baseline topology minus *Pan troglodytes*). The tree is rooted with *Colobus*. Purple diamonds indicate all changes in relation to the baseline, but only the differences in the hominin + *Pan* subtree topology were considered in the RF calculation.

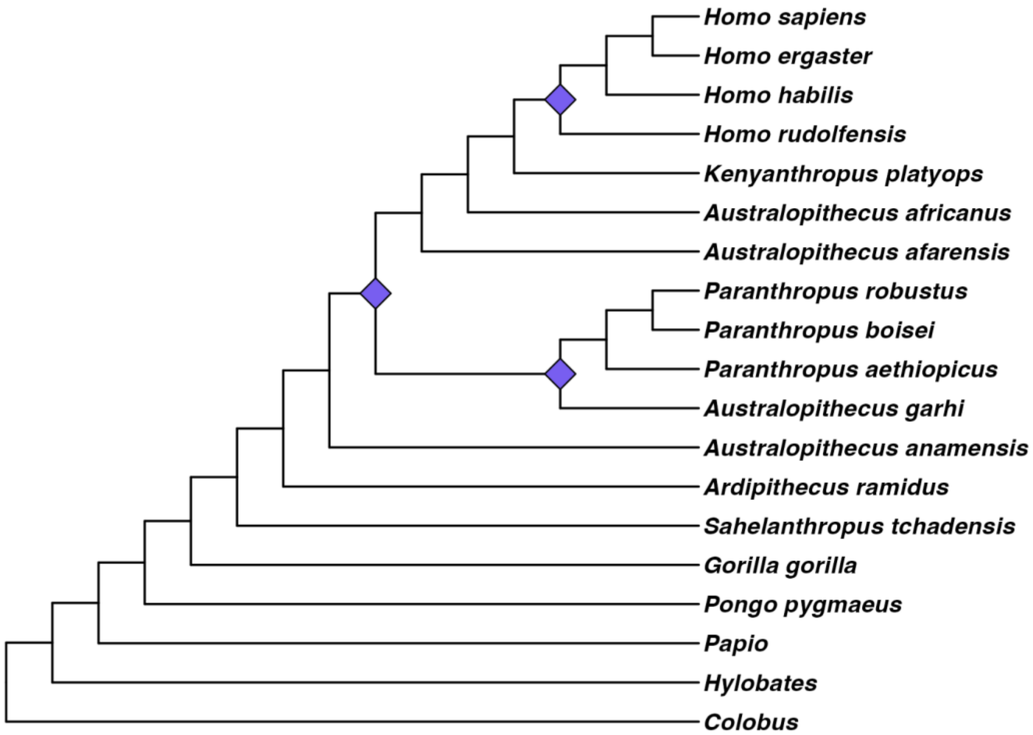

**Figure S10. Hominin phylogenetic tree topology inferred without *Au. anamensis*.** The consensus tree topology inferred from the character matrix excluding *Au. anamensis* using the branch and bound algorithm with maximum clade credibility ($\boldsymbol{RF = 8}$ between this and the baseline topology minus *Au. anamensis*). The tree is rooted with *Colobus*. Purple diamonds indicate all changes in relation to the baseline, but only the differences in the hominin + *Pan* subtree topology were considered in the RF calculation.

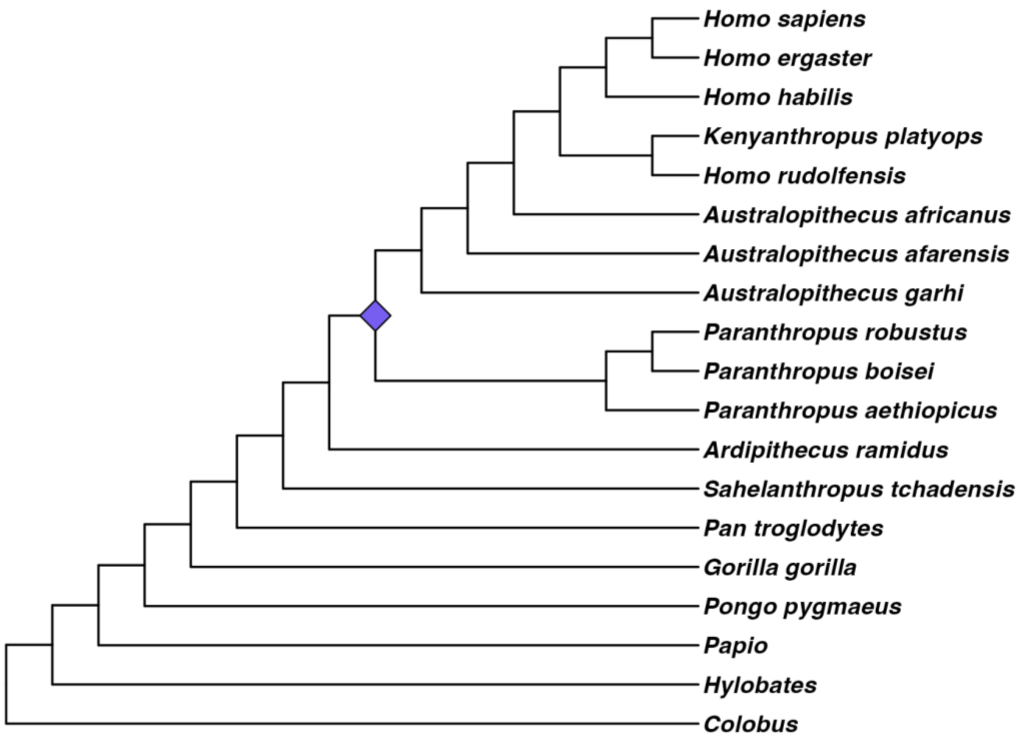

**Figure S11. Hominin phylogenetic tree topology inferred without *H. sapiens*.** The consensus tree topology inferred from the character matrix excluding *H. sapiens* using the branch and bound algorithm with maximum clade credibility ($\boldsymbol{RF = 8}$ between this and the baseline topology minus *H. sapiens*). The tree is rooted with *Colobus*. Purple diamonds indicate all changes in relation to the baseline, but only the differences in the hominin + *Pan* subtree topology were considered in the RF calculation.

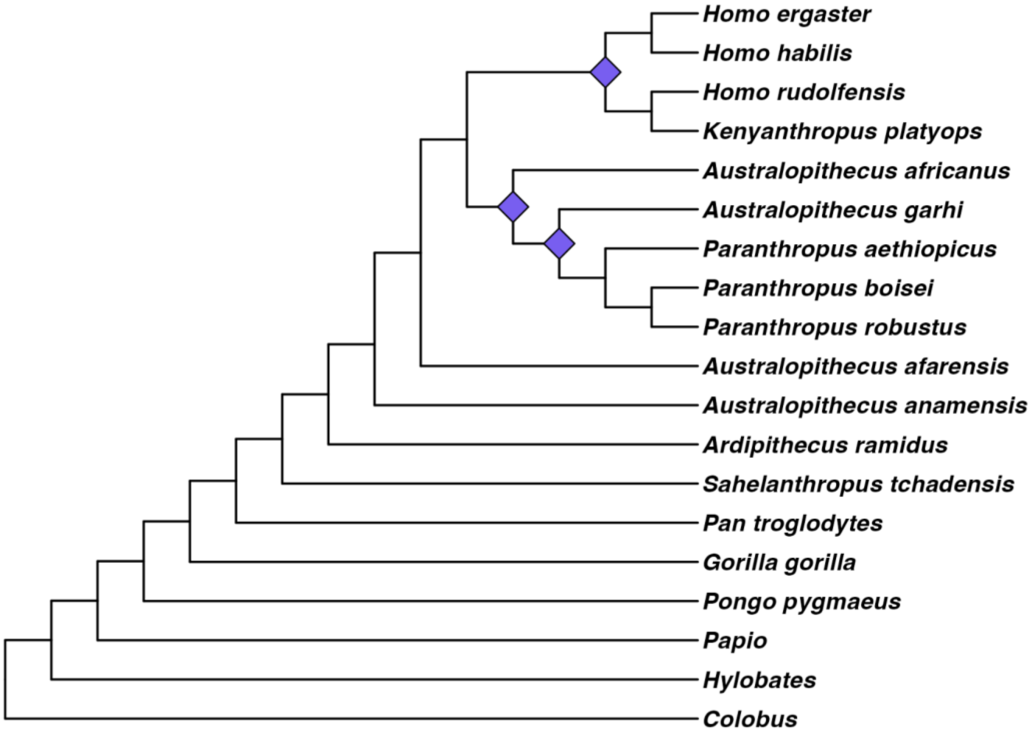

**Figure S12. Hominin phylogenetic tree topology inferred without *K. platyops*.** Consensus tree topology inferred from the character matrix excluding *K. platyops* using the branch and bound algorithm with maximum clade credibility ($RF = 8$ between this and the baseline topology minus *K. platyops*). The topology is rooted with *Colobus*. Purple diamonds show changes relative to the baseline topology.

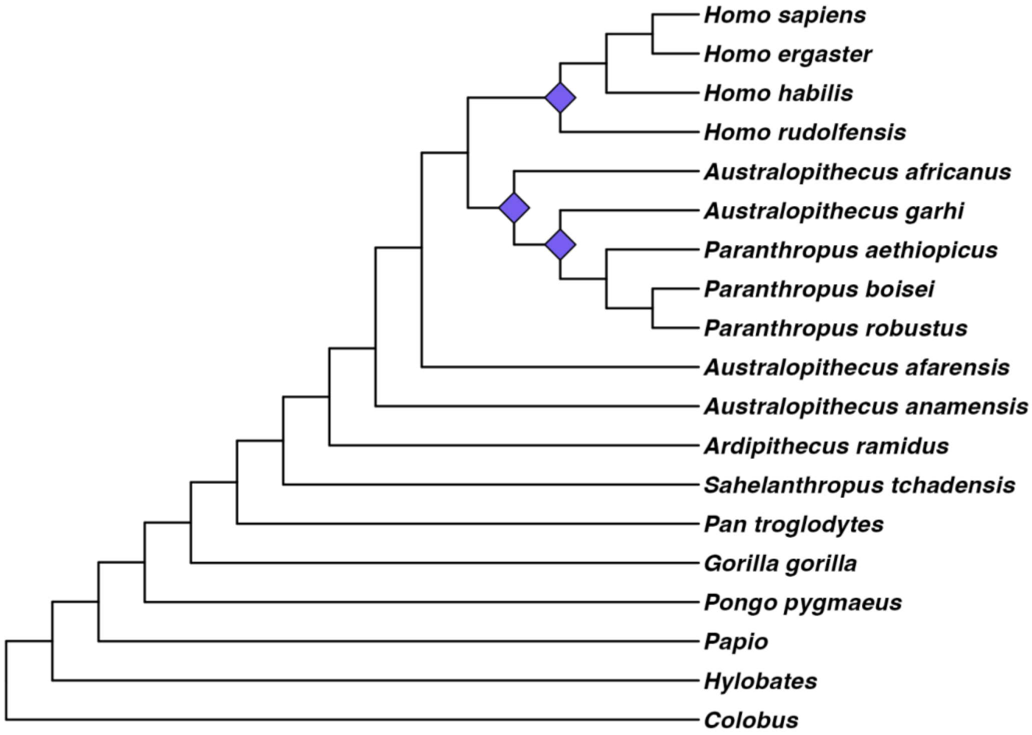


**Figure S13. Hominin phylogenetic tree topology inferred without *H. rudolfensis*.** The consensus tree topology inferred from the character matrix excluding *H. rudolfensis* using the branch and bound algorithm with maximum clade credibility ($RF = 8$ between this and the baseline topology minus *H. rudolfensis*). The tree is rooted with *Colobus*. Purple diamonds indicate all changes in relation to the baseline, but only the differences in the hominin + *Pan* subtree topology were considered in the RF calculation.

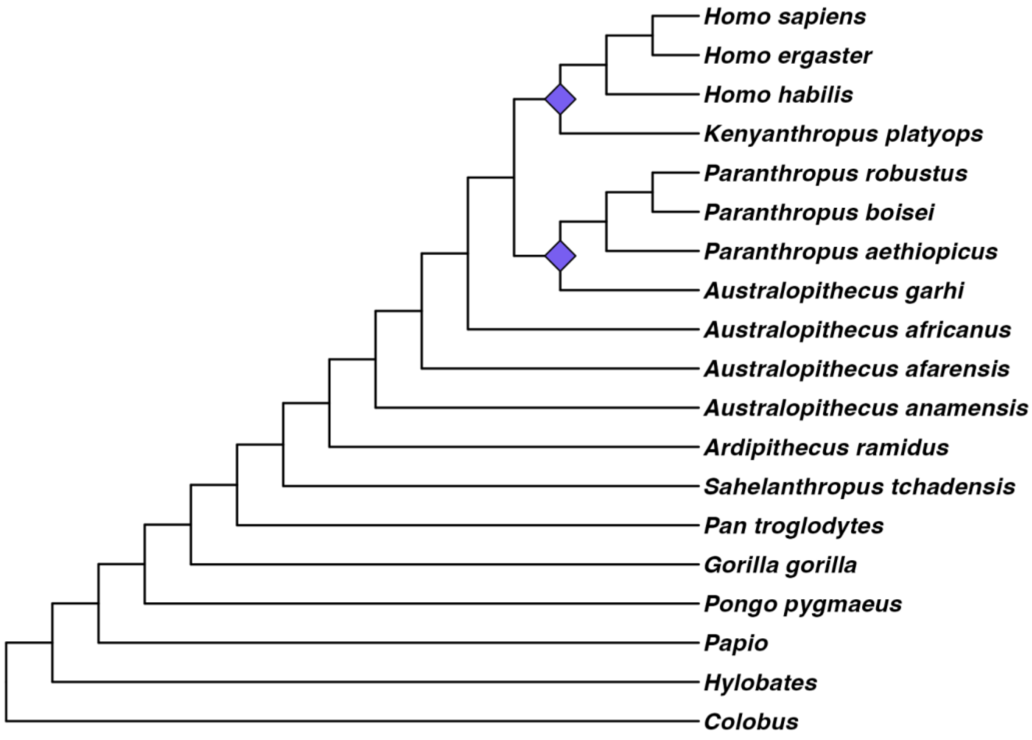
Homo sapiens
Homo ergaster
Homo habilis
Kenyanthropus platyops
Paranthropus robustus
Paranthropus boisei
Paranthropus aethiopicus
Australopithecus garhi
Australopithecus africanus
Australopithecus afarensis
Australopithecus anamensis
Ardipithecus ramidus
Sahelanthropus tchadensis
Pan troglodytes
Gorilla gorilla
Pongo pygmaeus
Papio
Hylobates
Colobus

**Figure S14. Hominin phylogenetic tree topology inferred without *P. robustus*.** The consensus tree topology inferred from the character matrix excluding *P. robustus* using the branch and bound algorithm with maximum clade credibility ($\boldsymbol{RF = 8}$ between this and the baseline topology minus *P. robustus*). The tree is rooted with *Colobus*. Purple diamonds indicate all changes in relation to the baseline, but only the differences in the hominin + *Pan* subtree topology were considered in the RF calculation.

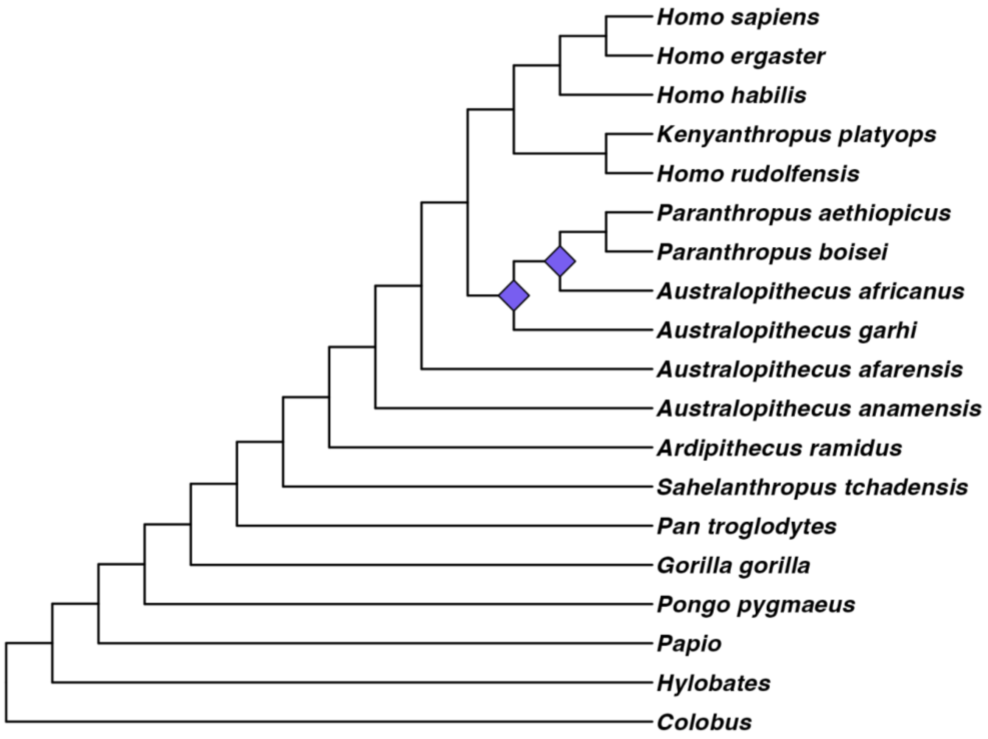

**Figure S15. Hominin phylogenetic tree topology inferred without *Au. afarensis*.** The consensus tree topology inferred from the character matrix excluding *Au. afarensis* using the branch and bound algorithm with maximum clade credibility ($\boldsymbol{RF = 6}$ between this and the baseline topology minus *Au. afarensis*). The tree is rooted with *Colobus*. Purple diamonds indicate all changes in relation to the baseline, but only the differences in the hominin + *Pan* subtree topology were considered in the RF calculation.

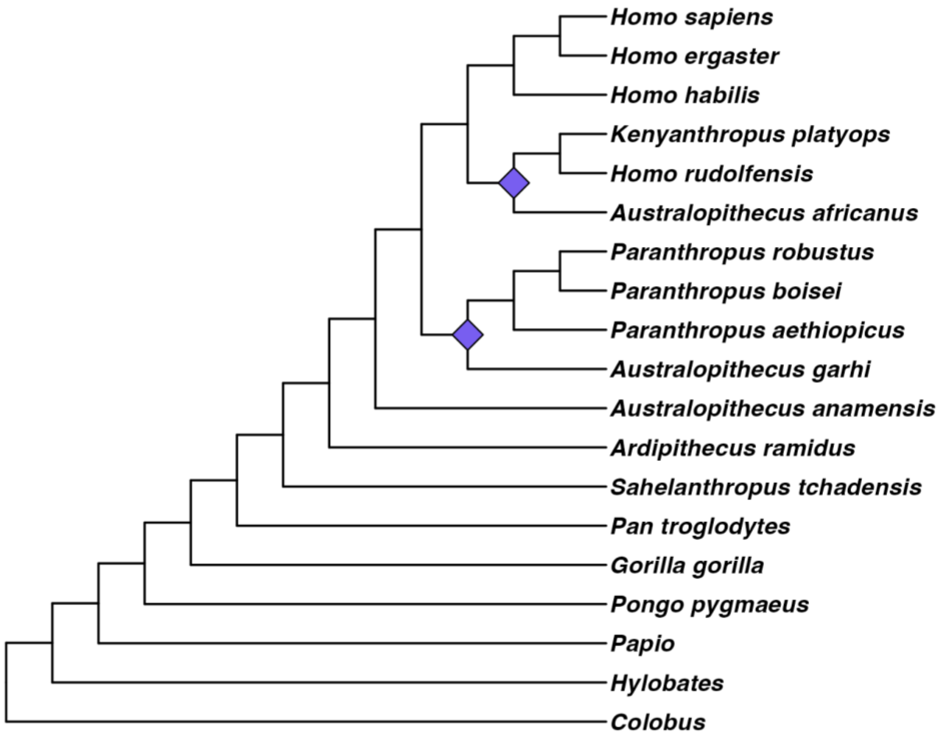

**Figure S16. Hominin phylogenetic tree topology inferred without *Au. africanus*.** The consensus tree topology inferred from the character matrix excluding *Au. africanus* using the branch and bound algorithm with maximum clade credibility ($\boldsymbol{RF = 6}$ between this and the baseline topology minus *Au. africanus*). The tree is rooted with *Colobus*. Purple diamonds indicate all changes in relation to the baseline, but only the differences in the hominin + *Pan* subtree topology were considered in the RF calculation.

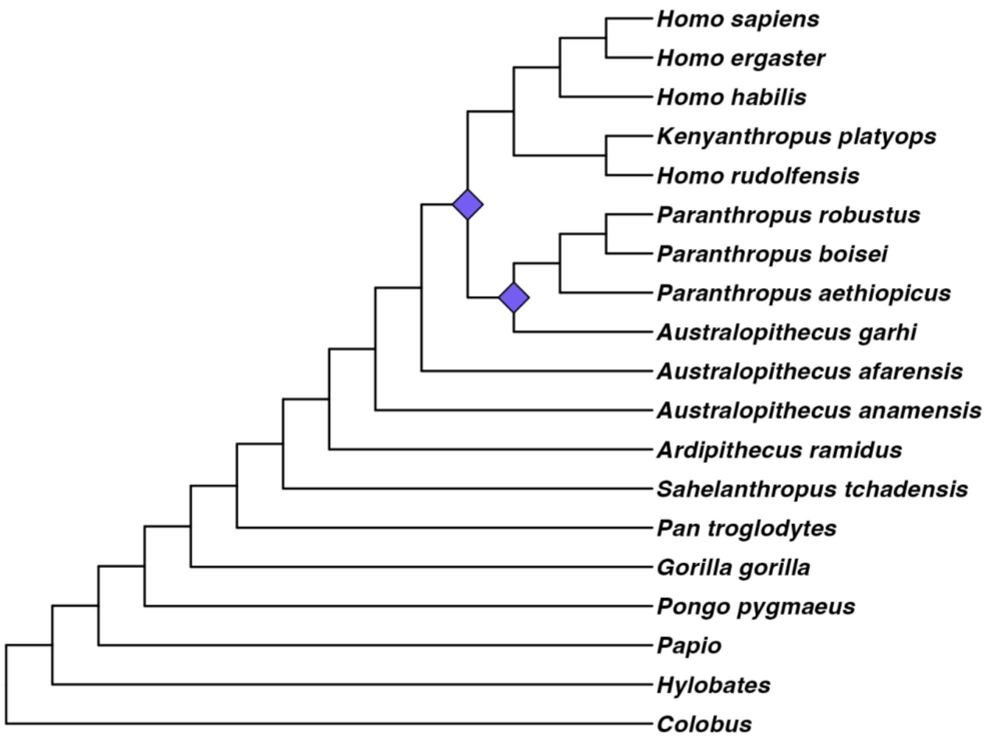

**Figure S17. Hominin phylogenetic tree topology inferred without *H. habilis*.** The consensus tree topology inferred from the character matrix excluding *H. habilis* using the branch and bound algorithm with maximum clade credibility ($\boldsymbol{RF = 6}$ between this and the baseline topology minus *H. habilis*). The tree is rooted with *Colobus*. Purple diamonds indicate all changes in relation to the baseline, but only the differences in the hominin + *Pan* subtree topology were considered in the RF calculation.

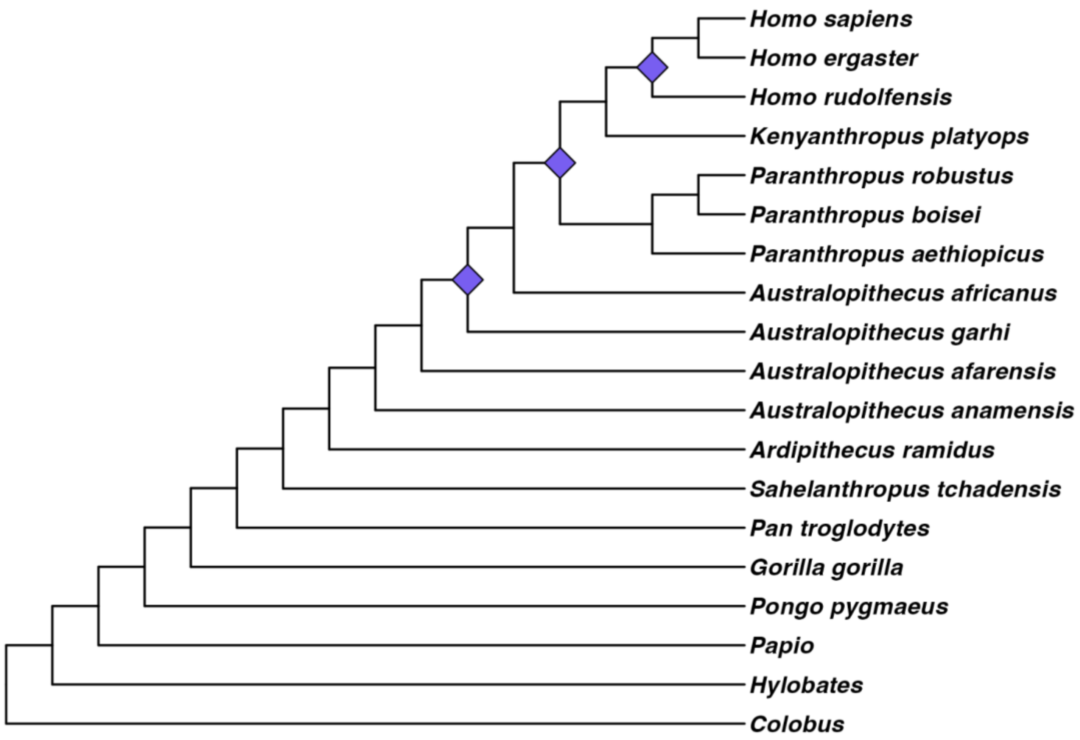

**Figure S18. Hominin phylogenetic tree topology inferred without *Colobus*.** The consensus tree topology inferred from the character matrix excluding *Colobus* using the branch and bound algorithm with maximum clade credibility ($\boldsymbol{RF = 4}$ between this and the baseline topology minus *Colobus*). The tree is rooted with *Papio*, the other Monkey of Africa and Eurasia included in this character matrix. Purple diamonds indicate all changes in relation to the baseline, but only the differences in the hominin + *Pan* subtree topology were considered in the RF calculation.

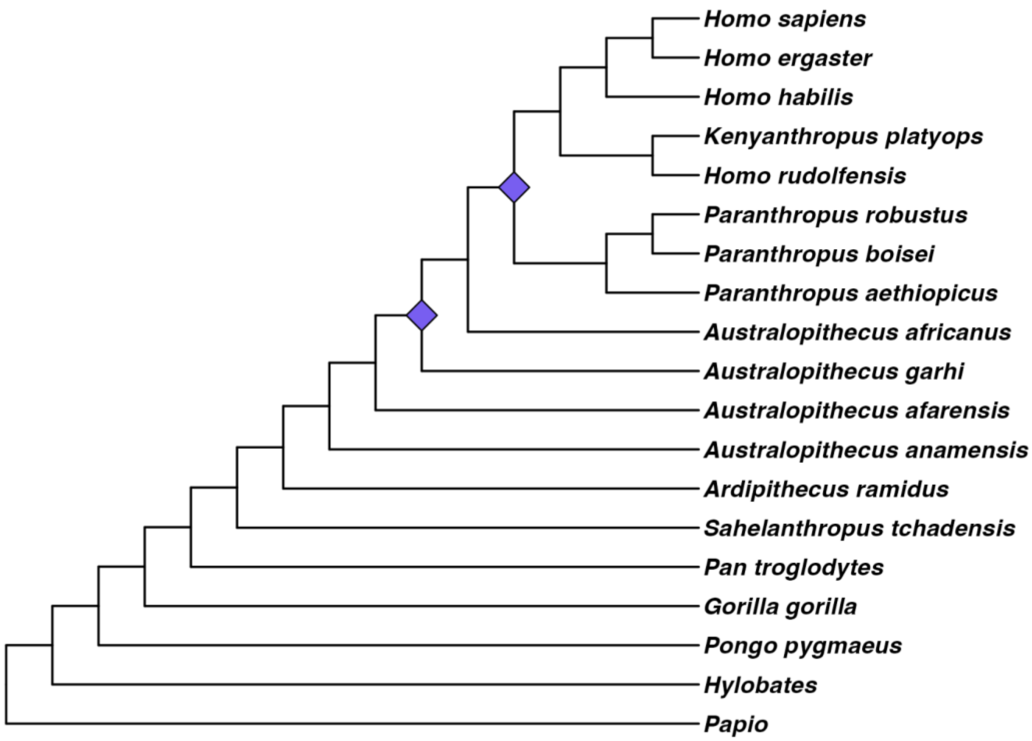

**Figure S19. Hominin phylogenetic tree topology inferred without *G. gorilla*.** The consensus tree topology inferred from the character matrix excluding *G. gorilla* using the branch and bound algorithm with maximum clade credibility ($\boldsymbol{RF = 4}$ between this and the baseline topology minus *G. gorilla*). The tree is rooted with *Colobus*. Purple diamonds indicate all changes in relation to the baseline, but only the differences in the hominin + *Pan* subtree topology were considered in the RF calculation.

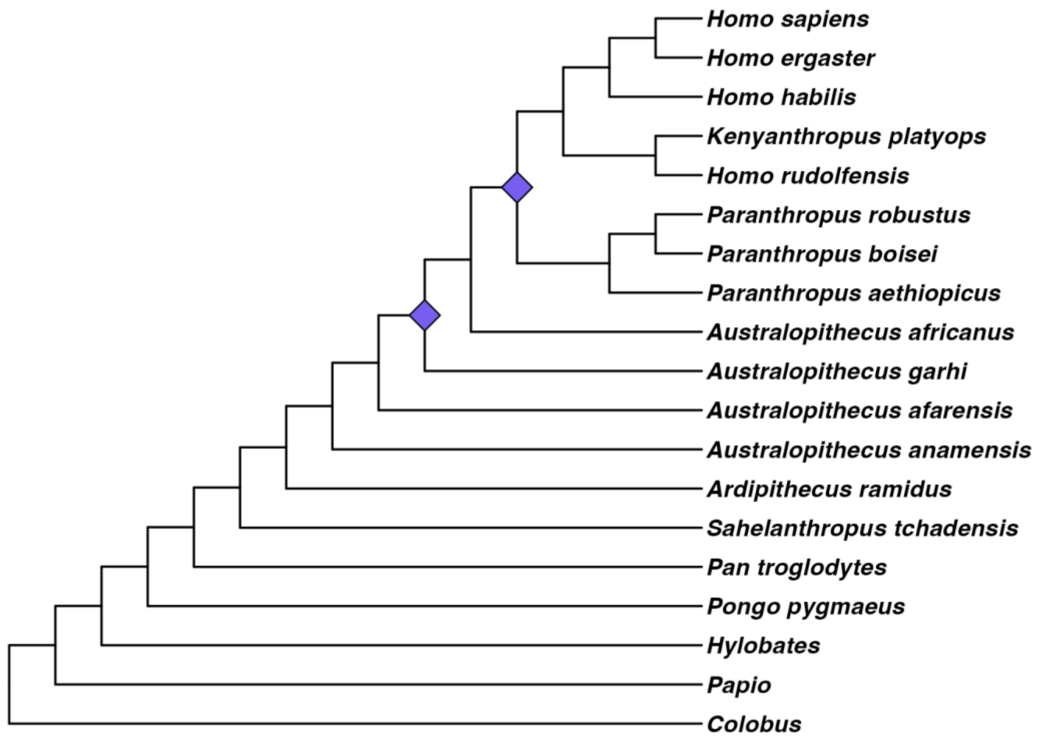

**Figure S20. Hominin phylogenetic tree topology inferred without *Hylobates*.** The consensus tree topology inferred from the character matrix excluding *Hylobates* using the branch and bound algorithm with maximum clade credibility ($\boldsymbol{RF = 4}$ between this and the baseline topology minus *Hylobates*). The tree is rooted with *Colobus*. Purple diamonds indicate all changes in relation to the baseline, but only the differences in the hominin + *Pan* subtree topology were considered in the RF calculation.

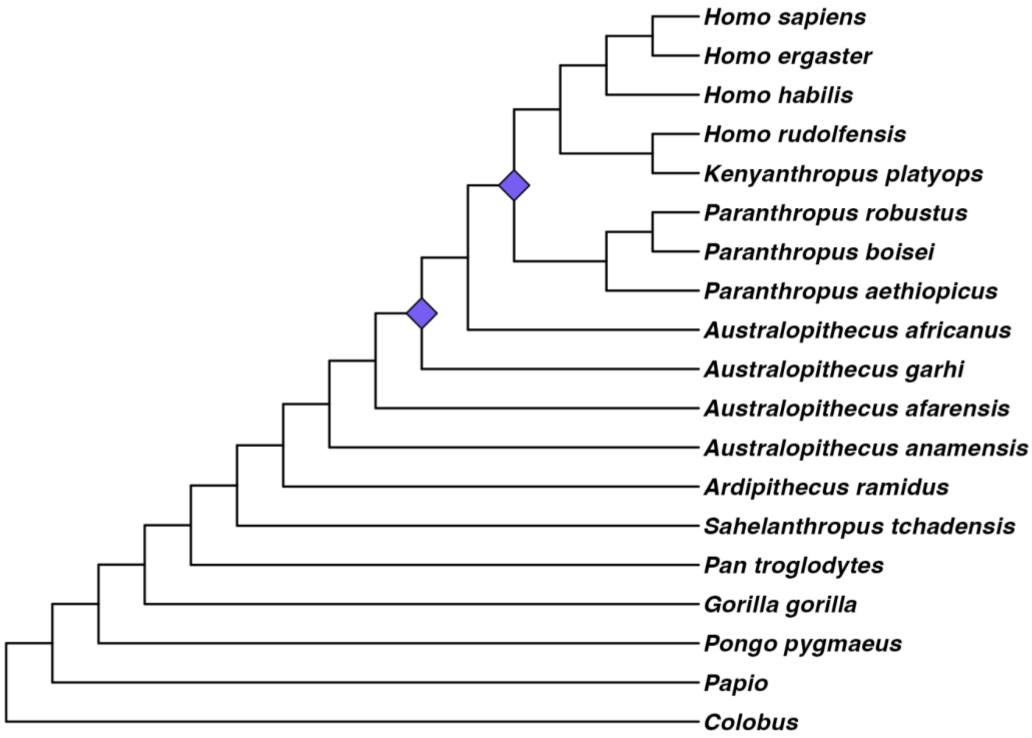

**Figure S21. Hominin phylogenetic tree topology inferred without *H. ergaster*.** The consensus tree topology inferred from the character matrix excluding *H. ergaster* using the branch and bound algorithm with maximum clade credibility ($\boldsymbol{RF = 4}$ between this and the baseline topology minus *H. ergaster*). The tree is rooted with *Colobus*. Purple diamonds indicate all changes in relation to the baseline, but only the differences in the hominin + *Pan* subtree topology were considered in the RF calculation.

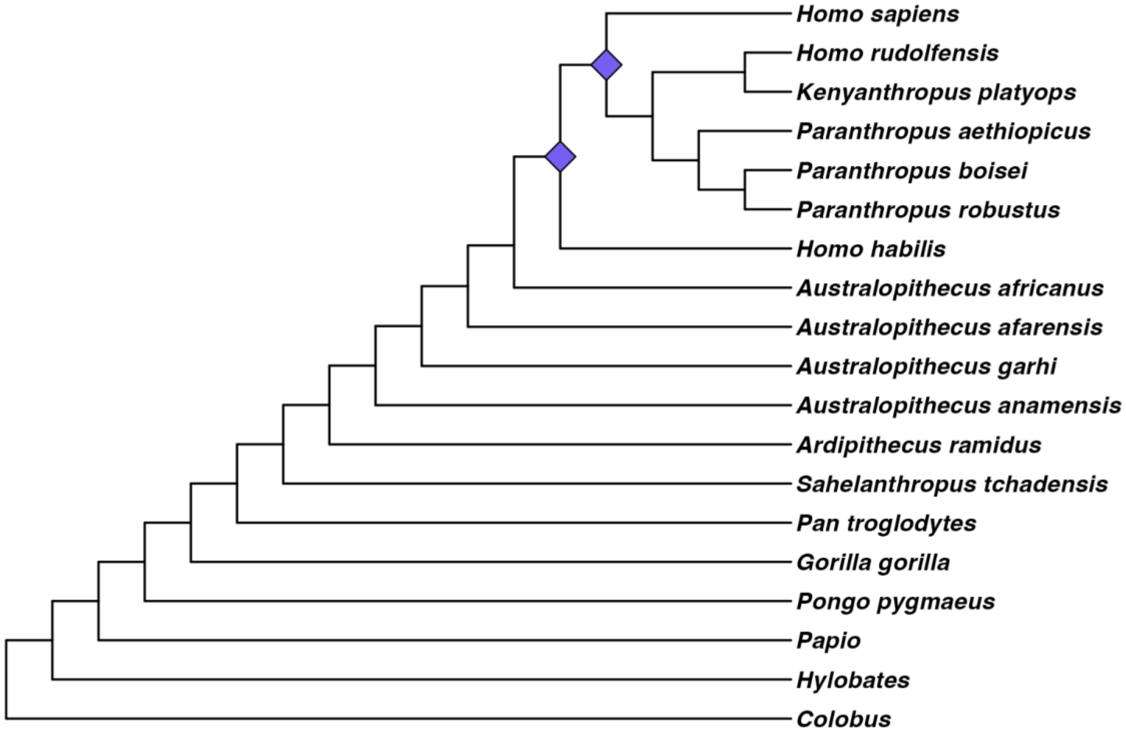

**Figure S22. Hominin phylogenetic tree topology inferred without *Papio*.** The consensus tree topology inferred from the character matrix excluding *Papio* using the branch and bound algorithm with maximum clade credibility ($RF = 4$ between this and the baseline topology minus *Papio*). The tree is rooted with *Colobus*. Purple diamonds indicate all changes in relation to the baseline, but only the differences in the hominin + *Pan* subtree topology were considered in the RF calculation.

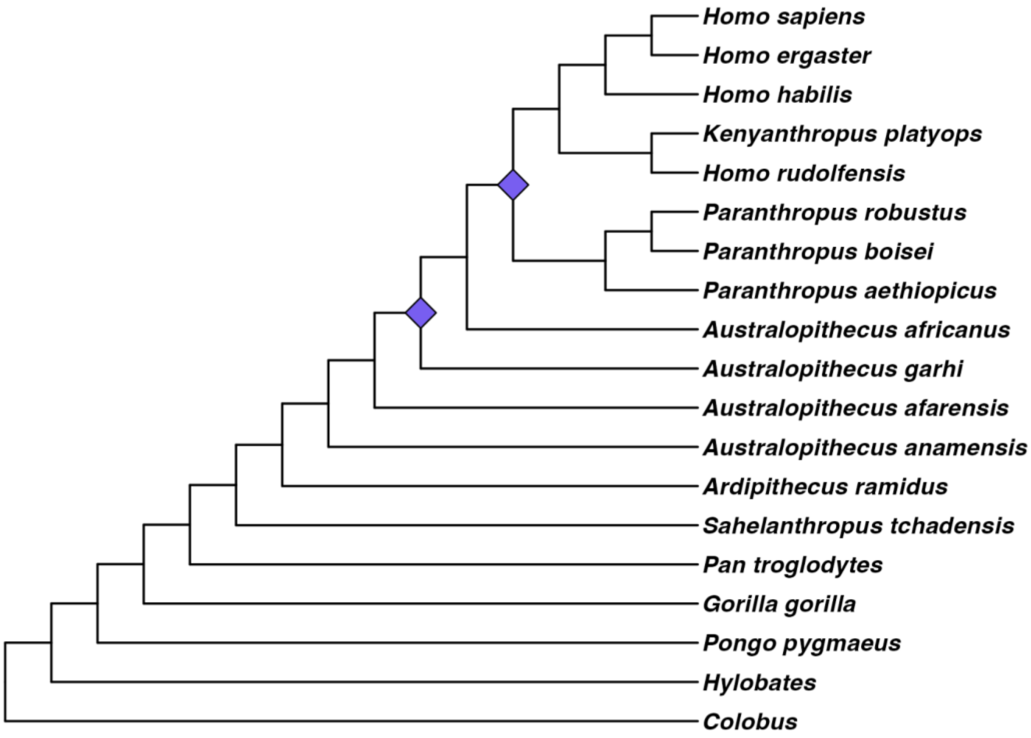

**Figure S23. Hominin phylogenetic tree topology inferred without *P. boisei*.** The consensus tree topology inferred from the character matrix excluding *P. boisei* using the branch and bound algorithm with maximum clade credibility ($\boldsymbol{RF = 4}$ between this and the baseline topology minus *P. boisei*). The tree is rooted with *Colobus*. Purple diamonds indicate all changes in relation to the baseline, but only the differences in the hominin + *Pan* subtree topology were considered in the RF calculation.

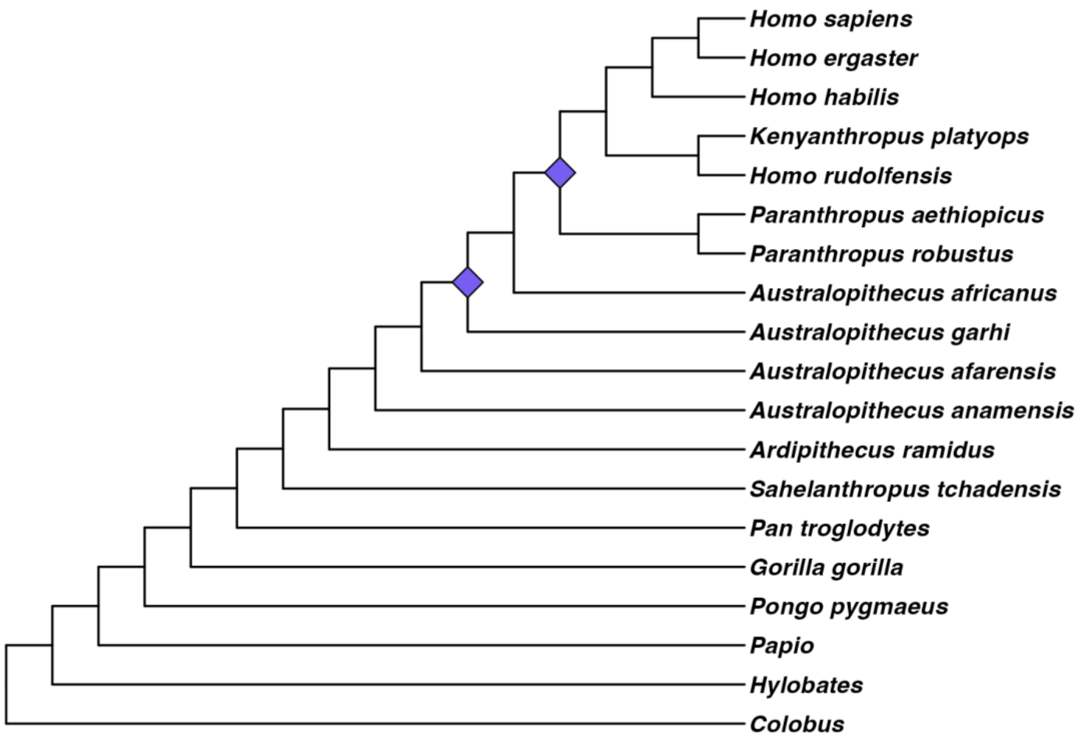

**Figure S24. Hominin phylogenetic tree topology inferred without *S. tchadensis*.** The consensus tree topology inferred from the character matrix excluding *S. tchadensis* using the branch and bound algorithm with maximum clade credibility ($\boldsymbol{RF = 4}$ between this and the baseline topology minus *S. tchadensis*). The tree is rooted with *Colobus*. Purple diamonds indicate all changes in relation to the baseline, but only the differences in the hominin + *Pan* subtree topology were considered in the RF calculation.

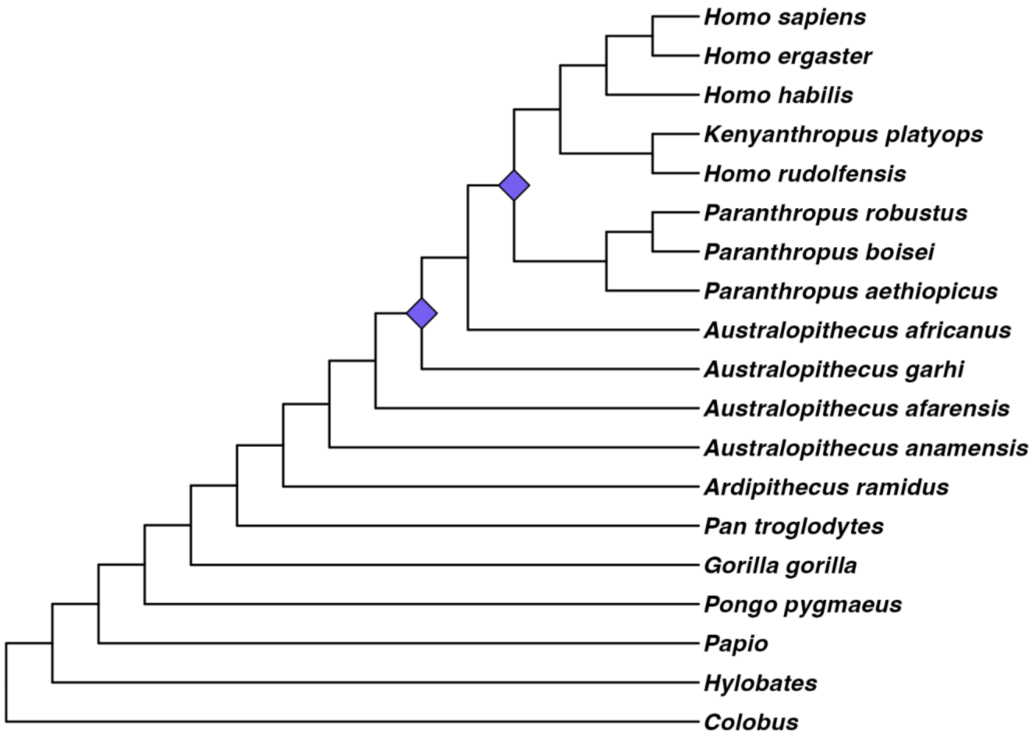
